# Generation of strong casein kinase 1 inhibitor of *Arabidopsis thaliana*

**DOI:** 10.1101/642884

**Authors:** Ami N Saito, Hiromi Matsuo, Keiko Kuwata, Azusa Ono, Toshinori Kinoshita, Junichiro Yamaguchi, Norihito Nakamichi

## Abstract

Casein kinase 1 (CK1) is an evolutionarily conserved protein kinase among eukaryotes. Studies on yeast, fungi, and animals have revealed that CK1 plays roles in divergent biological processes. By contrast, the collective knowledge regarding the biological roles of plant CK1 lags was behind those of animal CK1. One of reasons for this is that plants have more multiple genes encoding CK1 than animals. To accelerate the research for plant CK1, a strong CK1 inhibitor that efficiently inhibits multiple members of CK1 proteins in vivo (in planta) is required. Here, we report a novel strong CK1 inhibitor of Arabidopsis (AMI-331). Using a circadian period-lengthening activity as estimation of the CK1 inhibitor effect in vivo, we performed a structure-activity relationship (SAR) study of PHA767491 (1,5,6,7-tetrahydro-2-(4-pyridinyl)-4H-pyrrolo[3,2-c]pyridin-4-one hydrochloride), a potent CK1 inhibitor of Arabidopsis, and found that PHA767491 analogues bearing a propargyl group at the pyrrole nitrogen atom (AMI-212) or a bromine atom at the pyrrole C3 position (AMI-23) enhance the period-lengthening activity. The period lengthening activity of a hybrid molecule of AMI-212 and AMI-23 (AMI-331) is about 100-fold stronger than that of PHA767491. An in vitro assay indicated a strong inhibitory activity of CK1 kinase by AMI-331. Also, affinity proteomics using an AMI-331 probe showed that targets of AMI-331 are mostly CK1 proteins. As such, AMI-331 is a strong potent CK1 inhibitor that shows promise in the research of CK1 in plants.

## Introduction

Casein kinase 1 (CK1) is an evolutionarily conserved serine-threonine protein kinase among eukaryotes. CK1 plays key roles in various biological processes in these species. For instance, yeast CK1 isoforms are implicated in such as DNA repair, cytokinesis, cell cycle progression, and vesicular trafficking (Gross and Anderson, 1998). Fungi CK1 controls circadian clock (Gorl et al., 2001). Mammal CK1 proteins are involved in various biological processes such as immune responses, cell cycle progression, DNA damage signal transduction, circadian clock, apoptosis, and signaling pathways for development (Knippschild et al., 2014).

In plants, CK1 family also regulates various physiological processes. For example, CKL2 is involved in stomatal closure (Zhao et al., 2016), CKL3 and CKL4 regulate the blue light signaling pathway (Tan et al., 2013), CKL6 control cortical microtubules (Ben-Nissan et al., 2008), and CKL8 is implicated in ethylene production (Tan and Xue, 2014). In many cases, CKL phosphorylates substrate proteins, and phosphorylation by CKL triggers two distinct effects: degradation of substrates, or modification of substrate activity. Phosphorylation of CRY proteins by CKL3 and CKL4 is related to CRY degradation (Tan et al., 2013). CKL8 phosphorylates ACS5 protein for degradation control (Tan and Xue, 2014). On the other hand, CKL2 regulates F-actin disassembly of ADF4 by phosphorylation (Zhao et al., 2016). CKL6 controls microtubule dynamics by phosphorylating tubulin (Ben-Nissan et al., 2008). Rice HDB2 belongs to the CK1 family, and is involved in regulating reproductive isolation (or hybrid breakdown) (Yamamoto et al., 2010), and root development and hormone sensitivity (Liu et al., 2003), although substrates of rice CK1 have not been identified. It should be noted that genetic redundancy among multiple members of CKL (e.g., 13 CKLs in Arabidopsis) might make further finding of biological processes regulated by the CK1 family challenging.

Employing small molecules or inhibitors of CK1 is another way to examine whether CK1 is involved in interesting biological processes. For example, IC261 had mostly been used for this purpose (see structure in Figure 1A). Recent studies used PF-670462 (see Figure 1B), which is a more potent and specific inhibitor of plant CK1 (Mizoi et al., 2019) (Uehara et al., 2019). Chemical screening combined with target identification of the hit molecule indicated that PHA767491 (see Figure 1C), a mammal CDK7 (cyclin dependent kinase 7) inhibitor, also targets plant CK1 (Uehara et al., 2019). Combinational use of PF-6700462 and PHA767491 proved that CK1 is involved in Arabidopsis circadian clock (Uehara). However, the required concentration of these molecules to modulate biological processes (or give phenotypes) was around 100 μM. Reducing the effective concentration is required to enhance the utility of these molecules.

**Figure 1.**
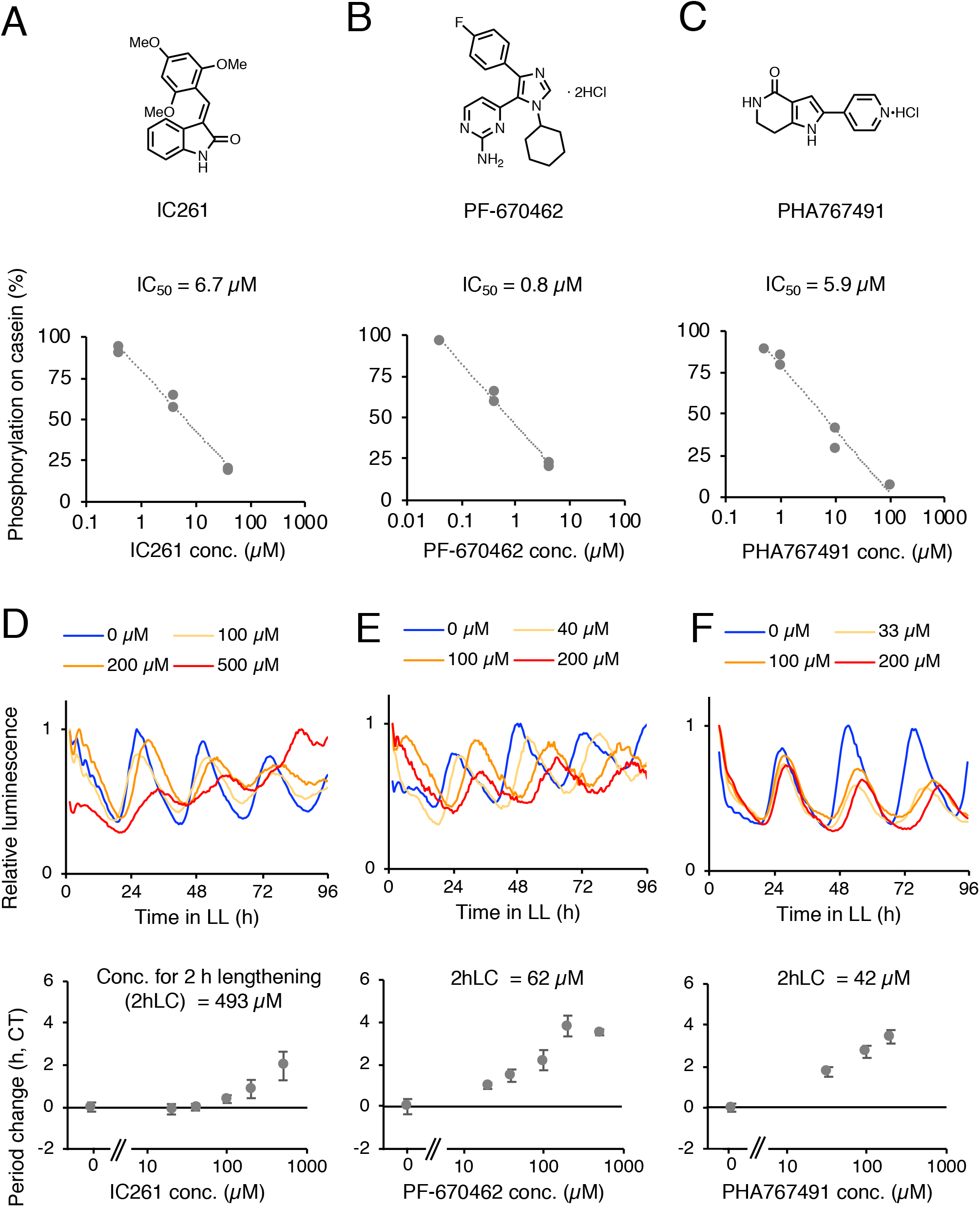
Activities of IC261, PF-670462, and PHA767491. Chemical structures of IC261 **(A)**, PF-670462 **(B)**, and PHA767491 **(C)**. In vitro CKL4 kinase activity with IC261 **(A)**, PF-670462 **(B)**, or PHA767491 **(C)**. Activity of circadian luciferase reporter *CCA1:LUC* with IC261 **(D)**, PF-670462 **(E)**, or PHA767491 **(F)**. Representative traces (upper) and increasing of period length relative to control experiments (lower). Error bars indicate SEM (n = 3∼12).

Through structure-activity relationship study of known plant CK1 inhibitor PHA767491, here we have developed a small molecule that has a strongest CK1 inhibitory activity in vivo. This molecule, named AMI-331, significantly lengthens the period of Arabidopsis circadian clock at concentrations below 1 μM. In addition, screening of proteins that were bound by AMI-331, indicated high specificity of AMI-331 on the CK1 family.

## Results

### Three CK1 inhibitors lengthen the circadian period

The activity of three known CK1 inhibitors was analyzed in vitro by a previously reported method (Uehara et al., 2019). Recombinant CKL4, model substrate casein, 32P-ATP, and different concentrations of the inhibitors were mixed in a reaction buffer and kept at 37°C for 2 h, and resulting samples were electrophoresed. 32P incorporation on casein was regarded as CKL4 kinase activity. IC_50_ (half maximal inhibitory concentration) was determined by the average of at least two independent experiments. IC_50_ of IC261, PF-670462, and PHA767491 were 6.7 μM, 0.8 μM, and 5.9 μM, respectively (Fig. 1A-C). A stronger in vitro CK1 inhibitory activity by PF-670462 over PHA767491 was consistent with our previous report (Uehara et al., 2019).

To estimate the CK1 inhibitor activity in vivo, we choose the circadian period lengthening activity. In addition, monitoring the circadian period using a clock reporter line (*CCA1:LUC*) and an automated luminometer render the evaluation of the molecules rapid. These tools can measure the circadian period with highly confidence, and to estimate in vivo CK1 inhibitor activity quantitatively (Uehara et al., 2019). Four day old *CCA1:LUC* seedlings grown under 12 h light / 12 h dark conditions were treated with IC261, PF-670462, and PHA767491 at different concentrations, and luminescence of seedlings were monitored under constant light conditions. Dimethyl sulfoxide (DMSO) treatment was used as a control experiment, since CK1 inhibitors were dissolved in DMSO (also see the methods section). IC261 lengthened the circadian period of *CCA1:LUC* seedlings in a dose-dependent manner (Fig. 1D). IC261 at 500 μM lengthened the period for 2 h. PF-670462 and PHA767491 also lengthened the period dose-dependently (Fig. 1E and F). Two-hour lengthening effects were found at about 90 μM PF-670472 and 47 μM PHA767491, which were consistent with our previous study (Uehara et al., 2019). Thus, PHA767491 was the strongest CK1 inhibitor in vivo among the three inhibitors, but the in vitro CK1 inhibitory activity of PHA767491 was weaker than that of PF-670462.

### PHA767491 derivatives modified on the pyrrole ring have strong period-lengthening activities

We sought to create a stronger CK1 inhibitor by modifying structure of PHA767491, since the in vitro CK1 inhibitor activity of PHA767471 was still weaker than that of PF-670472. We applied previous synthetic methods used for PHA767491 derivatives (Uehara et al., 2019) to make further derivatives of PHA767491. We firstly focused on modifying the pyrrole at the N-position, because two derivatives modified at this position retained period-lengthening activity (Uehara et al., 2019). The circadian period length of *CCA1:LUC* seedlings treated with newly synthesized PHA767491 derivatives was analyzed (Supplemental Figure 1). We found that derivatives bring a large substituent at the pyrrole N-position retained a weak period-lengthening activity compared to PHA767491 (AMI-113, 115, 116, 117, 118, 121, 122, 123, 130, 131, 132, 133, 134, and 214). Other types of derivatives with one or two chlorine atoms on the pyridine unit, possessed weaker activity as well (AMI-138 and 125). On the other hand, derivatives with ethyl group at pyrrole N-position possessed strong activity (AMI-126). Then, we tested whether further derivatization of AMI-126 with small alkyl groups can enhance activity. Among these derivatives, we found that AMI-212 has a stronger period-lengthening activity, compared to AMI-126. Further analysis showed that concentrations around 7.0 μM of AMI-212 lengthened the circadian period for 2 h, and 26 μM of AMI-212 lengthened the period for 5 h, showing about five-fold stronger activity compared to PHA767491 (Figure 2A).

**Figure 2.**
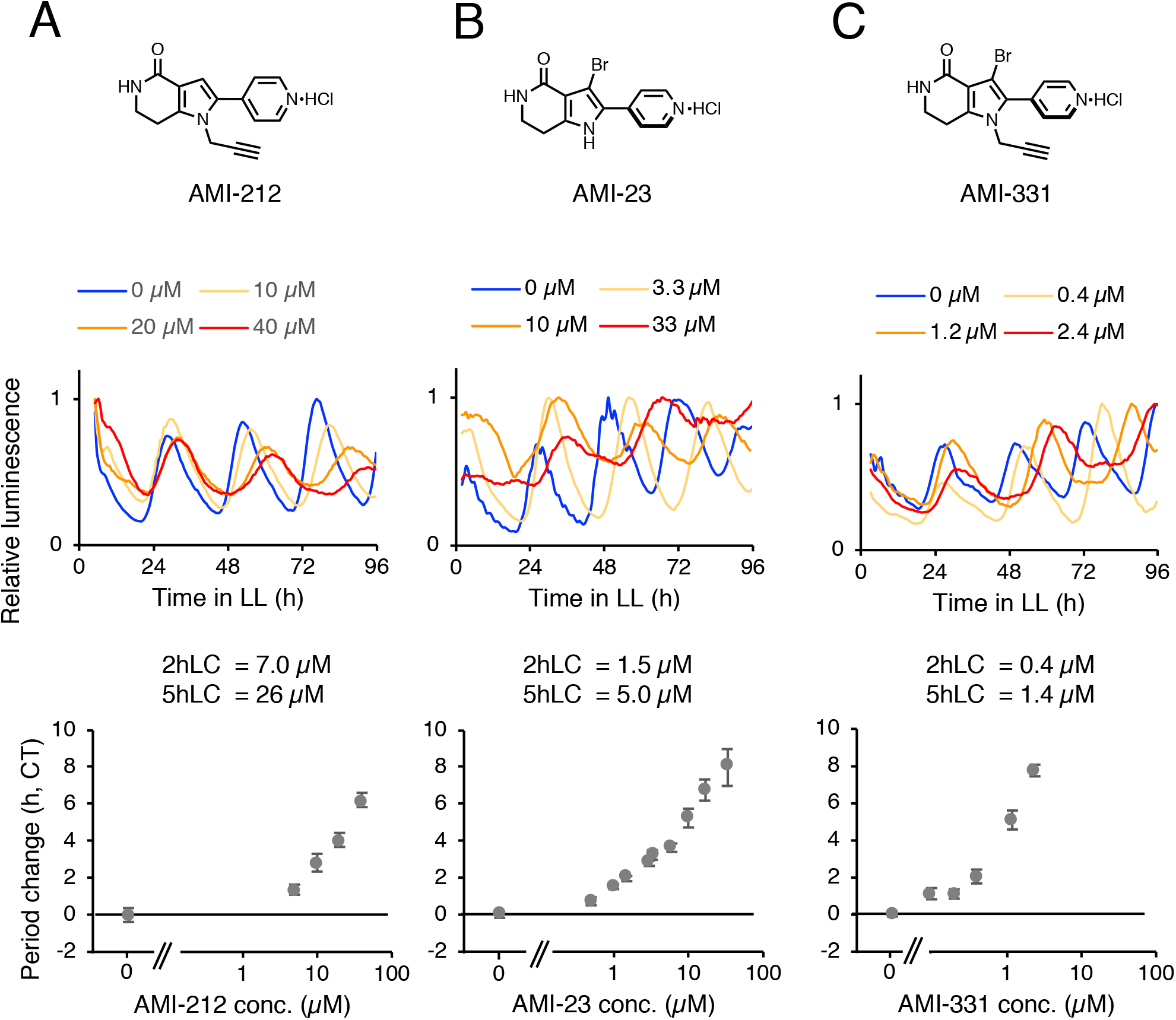
Period lengthening activity of AMI-212, AMI-23, and AMI-311. Structure of AMI-212 **(A)**, AMI-23 **(B)**, and AMI-331 **(C)** (upper). *CCA:LUC* activity with AMI-212 **(A)**, AMI-23 **(B)**, or AMI-331 **(C)** (middle and lower). Middle panels show representative traces, and lower panels show increasing of period length relative to control (n = 8 ∼22, with SEM).

Moreover, we found strong activities in three un-expected derivatives, which had a bromo-, chloro- or methyl-group at the pyrrole C3 position (AMI-23, 24, and 217, see Supplemental Figure 1). A strong period-lengthening activity of AMI-23 was found (Figure 2B). Concentrations of 1.5 μM and 5.0 μM of AMI-23 could lengthen the period for 2 h and 5 h, respectively.

We next examined the activity of the hybrid molecule of AMI-212 and AMI-23, namely AMI-331. Concentrations of 0.4 μM and 1.4 μM of AMI-331 lengthened the period for 2 h and 5 h, respectively (Figure 2C). Activity of AMI-331 was stronger than AMI-212 and AMI-23, and this was about 100-fold stronger than PHA767491.

### AMI-331 causes hyper accumulation of two clock-associated transcription factors

To test whether the period lengthening activity of AMI-331 is related to the inhibition of CK1 in vivo, we analyzed the amount of PRR5 in plants treated with AMI-331 because treatment of PHA767491 results in hyper accumulation of PRR5 in vivo (Uehara et al., 2019). Seedlings expressing PRR5-FLAG under Cauliflower Mosaic Virus 35S promoter (*35Spro:PRR5-FLAG*) were grown under constant light conditions for 4 days, and treated with AMI-331 and kept under constant dark for 1 day. PRR5-FALG protein amounts were decreased under dark compared to light conditions, due to ZTL-dependent degradation as described in previous studies (Kiba et al., 2007; Fujiwara et al., 2008) (Uehara et al., 2019). AMI-331 treatment resulted in hyper accumulation of PRR5-FALG under dark (Figure 3A). We next examined whether AMI-331 cause accumulation of PRR5-VP in *35Spro:PRR5-VP*, which has an opposite phenotype of *35Spro:PRR5-FLAG*. This analysis led us to examine PRR5 accumulation by AMI-331 in plants having different clock phenotypes. AMI-331 caused accumulation of PRR5-VP (Figure 3B). These results indicated that PRR5 protein is accumulated by AMI-331, likely due to inhibition of CK1 activity that causes PRR5 degradation in vivo. Since PHA767491 treatment also attenuates degradation of TOC1 under dark (Uehara et al., 2019), we tested whether AMI-331 affects TOC1 amounts in vivo. TOC1 amounts were increased by AMI-331 treatments (Fig. 3). The effective concentrations of AMI-331 for PRR5 or TOC1 accumulations (10 to 50 μM) were far less than that of PHA767491 (250 to 500 μM) (Uehara et al., 2019), indicating a stronger activity for AMI-331 than PHA767491 in vivo.

**Figure 3.**
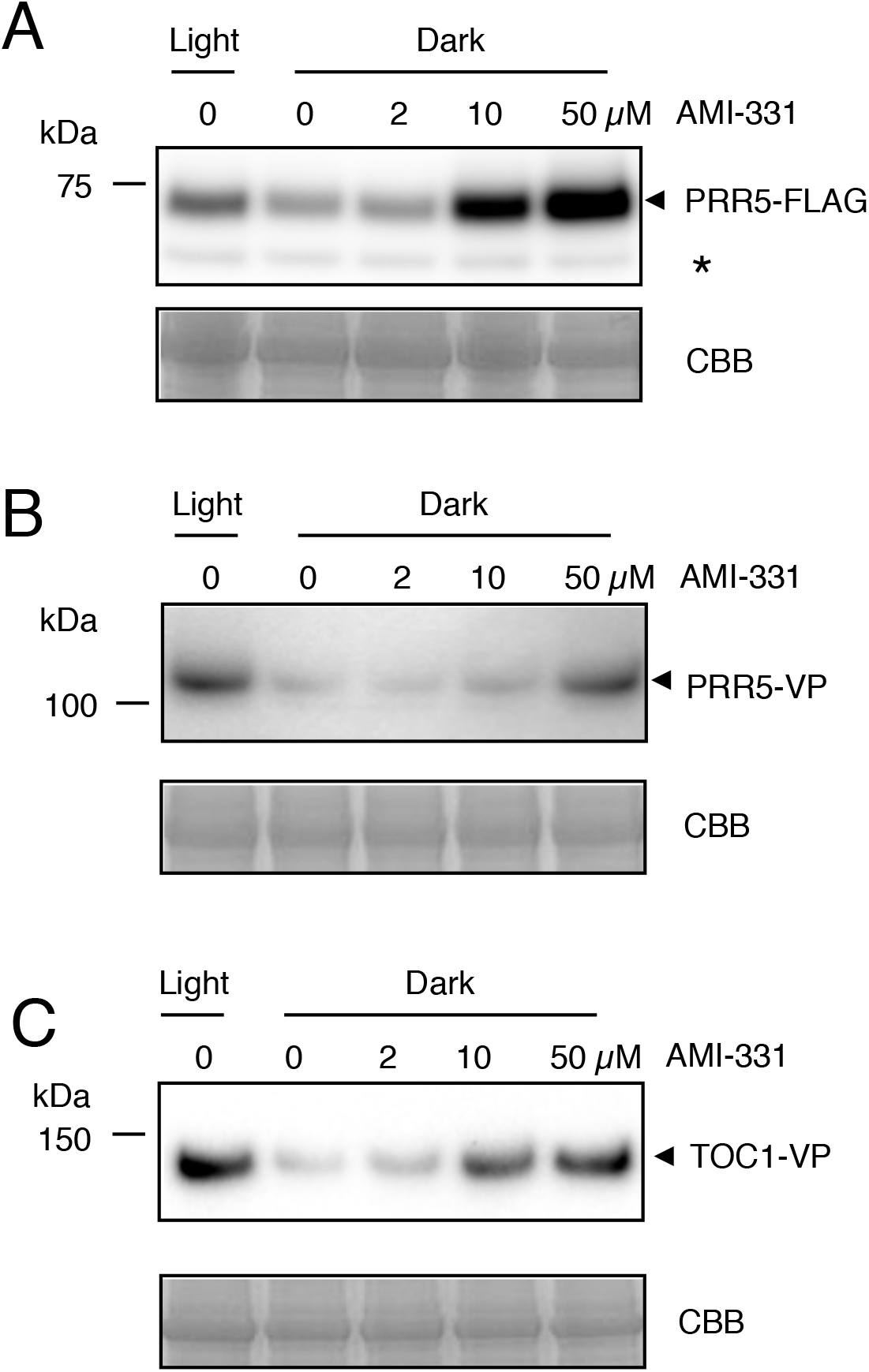
PRR5 and TOC1 proteins amount in plants treated with AMI-331. Protein amounts of PRR5-FLAG **(A)**, PRR5-VP **(B)**, and TOC1-VP **(C)** in the corresponding transgenic plants treated with AMI-331 (upper). Similar result was obtained in another trail (see supplemental figure 2).

### AMI-331 has a strong CK1 inhibitory activity in vitro

CK1 inhibitory activities of AMI-212, AMI-23, and AMI-331 in vitro were examined. IC_50_ values of CKL4 kinase activity by AMI-212, AMI-23, and AMI-331 were 1.2, 0.7, and 0.7, lower than that of PHA767491 (Fig. 4). This result suggested that strong in vitro CK1 inhibitory activity of AMI-331 causes strong CK1 inhibitory activity in vivo (period-lengthening activity and accumulation of PRR5 and TOC1). However, a very strong period-lengthening activity by AMI-331 compared to PHA767491 (∼100 times), as well as PRR5 accumulation activity (∼50 times), may suggests that unknown properties of AMI-331 contribute to strong in vivo strong period lengthening activity.

**Figure 4.**
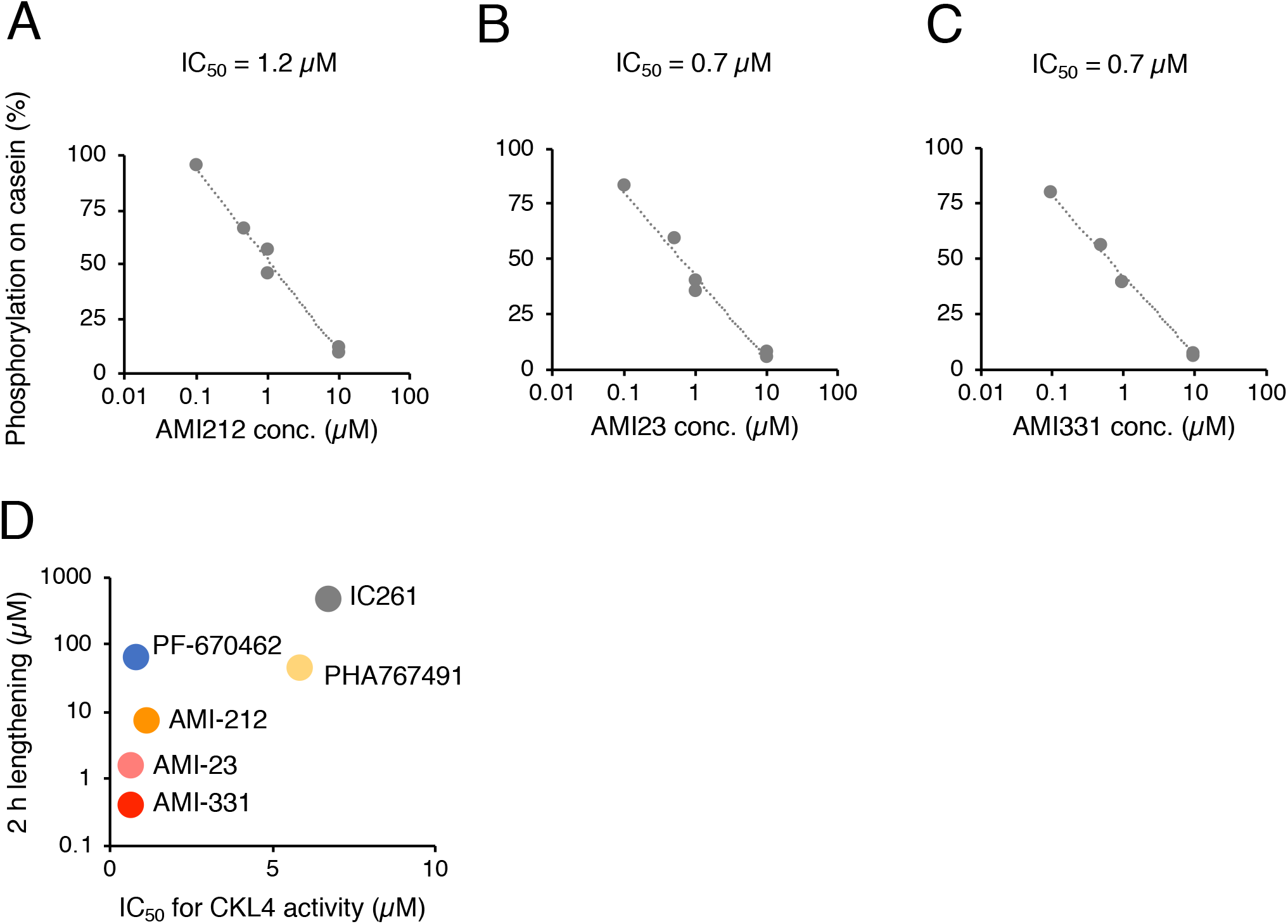
In vitro CKL4 kinase activity with three AMI molecules. In vitro CKL4 kinase activity with AMI-212 **(A)**, AMI-23 **(B)**, or AMI-331 **(C)**. **(D)** Comparisons of in vitro CK4 kinase activity (x-axis) and in vivo period lengthening activity (y-axis) with the CK1 inhibitors.

### Target identification of AMI-311

Strong in vitro CK1 inhibitory activity and in vivo activity related to the clock by AMI-331 suggested that AMI-331 modulates the clock via inhibition of CK1. However, one possible reason for strong biological activity of AMI-331 in vivo is that AMI-331 targets unexpected other proteins beside the CK1 family for period lengthening. To test this possibility, we set out to synthesize molecular probes of AMI-331 and perform screening direct target proteins using the probe. Since a previous study demonstrated that the nitrogen atom of the pyrrole ring of PHA767491 retains period-lengthening activity (Uehara et al., 2019), AMI-331 was also substituted with an alkyl group (AMI-329). AMI-329 retained weak but significant period lengthening activity (Fig. 5A). Then, AMI-329 was covalently bound to agarose beads, and the resulting AMI-329 beads were mixed with protein lysate of Arabidopsis seedlings, with or without AMI-331 (0, 5 or 50 μM) as competitor (Fig. 5B). Proteins whose digested peptides’ spectra over 2 in 0 μM AMI-331 sample were selected to ensure data integrity. In the reliable group, we further selected proteins whose relative spectra (spectra in 0 μM / summed spectra in 5 and 50 μM) were over 10, as AMI-331 bound proteins. The criteria contained 23 proteins, in which all members of the CK1 family and other 9 proteins were included (Fig. 5C). Spectra of these proteins were 0 to 7% in 5 μM AMI-331 sample, compared to sample without competitor (0 μM). These proteins spectra were zero in the sample containing 50 μM AMI-331. We also found other proteins were enriched in 0 μM sample compared to 5 and 50 μM samples. However, spectra numbers of these proteins were lesser than those of CKL family. In addition, spectra of reductase C (AT2G41680) and unknown protein (AT5G42765) in input fraction were more than those in 0 μM sample. Collectively, the analysis suggested that AMI-331 targets CKL family best, but also targets HYD1, LUP1, and YAK1.

**Figure 5.**
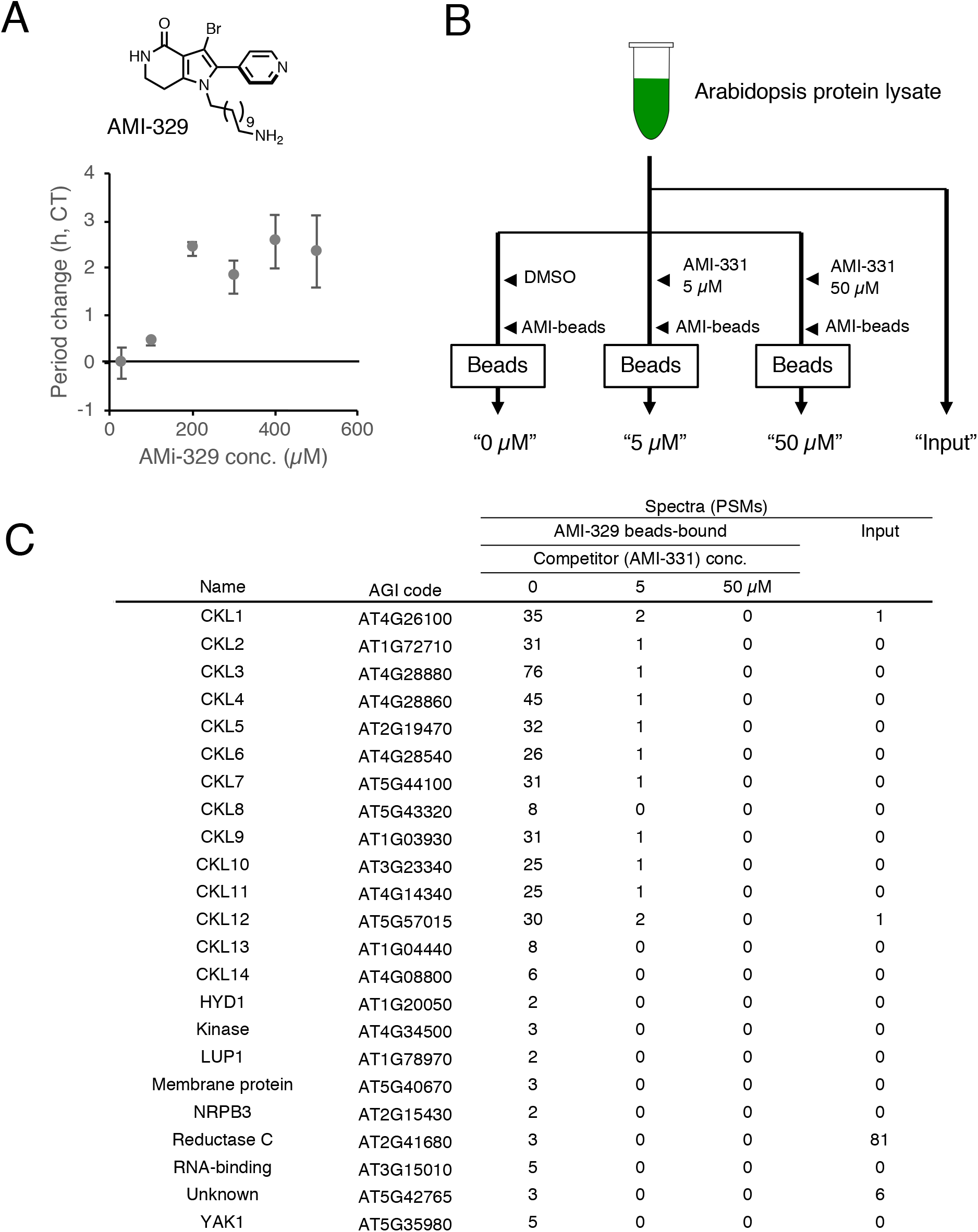
Target identification of AMI-331. **(A)** Structure and period lengthening activity of AMI-329. **(B)** Procedure of target identification of AMI-331. Targets of AMI-331 should be enriched in ‘0 μM’ compared to ‘5 μM’, ‘50 μM’, since free AMI-331 and AMI-329 beads competitively bind to AMI-331 targets. **(C)** Proteins that bound by AMI-329-beads. Spectra corresponding to the protein were shown. Note that two technical replicates were examined with similar results, so that these data were combined.

To access other potential targets of AMI-331, we analyzed spectra of proteins that were reported as PHA767491 targets (Uehara et al., PNAS 2019). Spectra of ATSK (GSK3) family were 2 to 31 in 0 μM AMI-331 sample (Supplemental Figure 3). These spectra were decreased to 14 to 53% in the 5 μM sample, and 0 to 13% in the 50 μM sample (Fig.6). Except for MPK5, spectra of MPK family were reliable (3 to 52) in the 0 μM sample. These spectra were decreased to 13 to 85% in 5 μM, and 0 to 31% in the 50 μM samples. Although CPK6 and CPK26 were reported as possible targets of PHA767491, all the CPK family was not enriched in the 0 μM sample compared to the 5 or 50 μM samples, suggesting that CPK is not a target of AMI-331. We further found that three other proteins encoding protein kinase (AT2G32850, AT3G58640, and AT3G61160) were also enriched in the 0 μM sample, compared to the 5 or 50 μM samples, suggesting that these proteins were possibly targets of AMI-331. The other proteins reported as possible targets of PHA767491 (who) were not enriched in the 0 μM AMI-331 sample, compared to the 5 or 50 μM samples.

**Figure 6.**
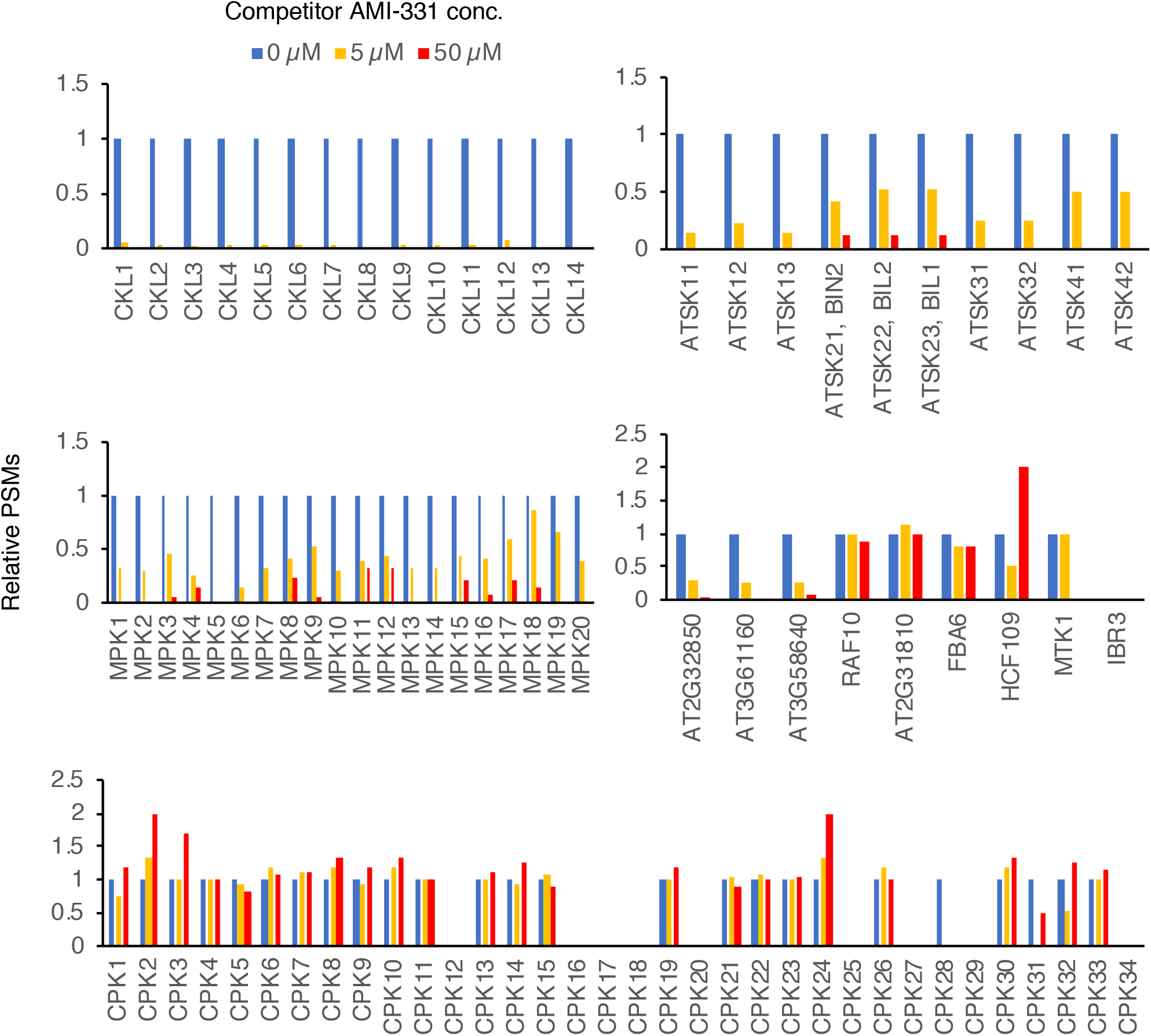
Relative spectra of potential PHA767491 target proteins in AMI-329-beads binding. Relative spectra to ‘0 μM’ were plotted. Actual spectra values were shown in supplemental figure 3.

These data again showed that AMI-331 prefers to target the CK1 family best, and AMI-331 may also targets the ATSK family, the MPK family, and other protein kinase (LUP, YAK1,, AT2G32850, AT3G58640, and AT3G61160).

### Effect of AMI-311 on CK1 and ATSK downstream genes expression

We next tried to understand the in vivo AMI-331 activity on other proteins beside the CK1 family. Since ATSK family was considered as potential targets of AMI-331, we analyzed whether AMI-331 treatment affects expression of genes downstream of ATSKs. To evaluate this approach, we used an ATSK-inhibitor, Bikinin (De Rybel et al., 2009), as a control experiment. Four-day-old Arabidopsis seedlings grown under constant light conditions were treated with Bikinin or AMI-331 for 6 h. RNA extracted from seedlings samples were analyzed by reverse transcription quantitative PCR analyses (RT-qPCR).

Treatment of both 10 and 50 μM samples of Bikinin resulted in reduction of expression of *GR60ox2* and *CPD* genes downstream of ATSK, as described previously (Fig. 7A) (De Rybel et al., 2009). Bikinin at 10 and 50 μM did not affect expression of *PRR7*, a downstream gene of the CK1 family. The result suggested that Bikinin perennially targets ATSK but not CK1.

**Figure 7.**
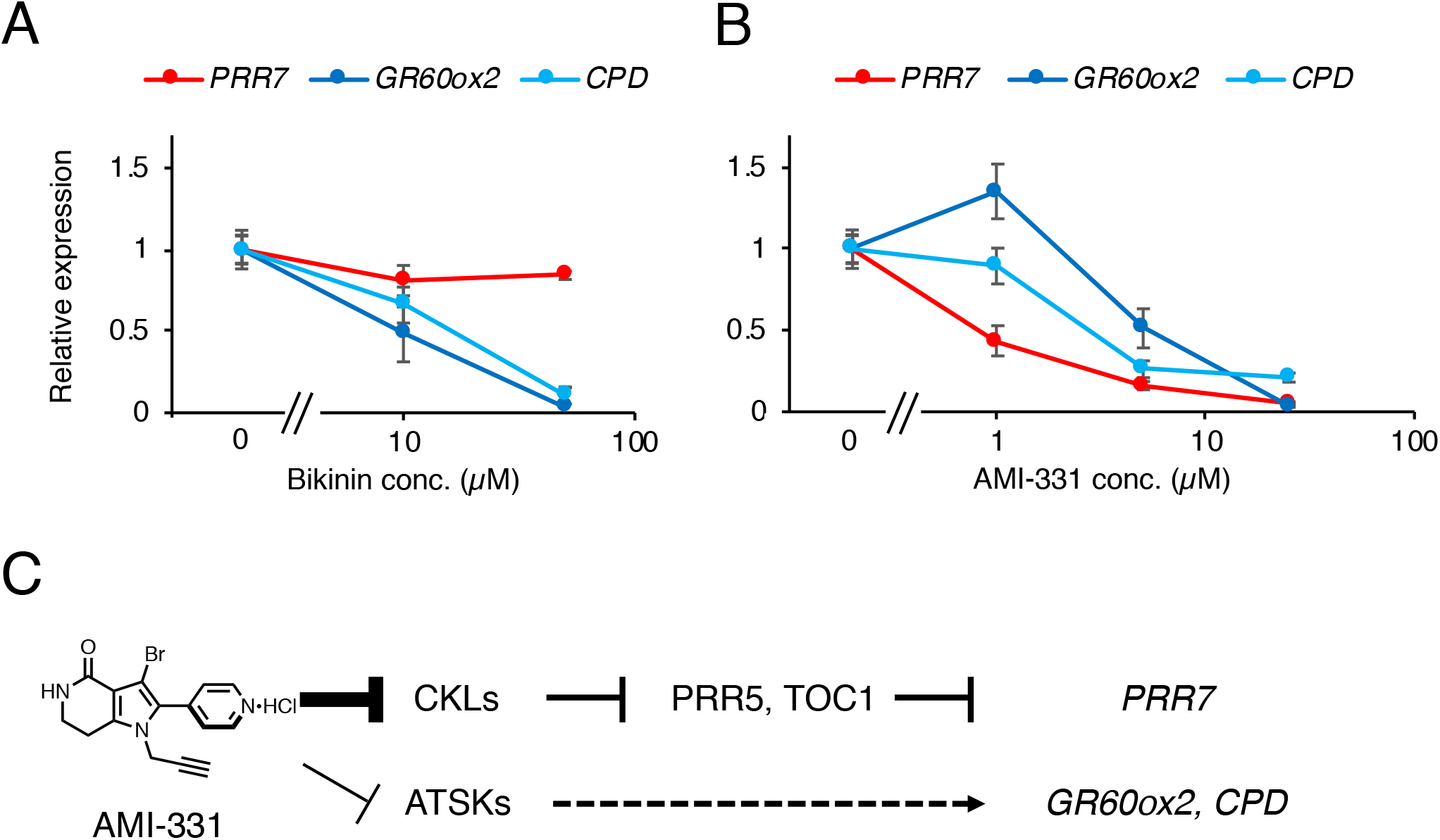
Expression of *PRR7, GR60ox2*, and *CPD* genes in plants treated with AMI-331. **(A)** Plants were treated with DMSO, 10 μM or 50 μM of Bikinin for 6 h, and gene expression were analyzed. **(B)** Plants were treated with DMSO, 1 μM, 5 μM, or 20 μM of AMI-331. Error bar indicates SEM of three biological replicates. Note that same DMSO samples were used for **(A)** and **(B)**. **(C)** AMI-331 activity for clock-related gene *PRR7* and brassinosteroid-related genes *GR60ox* and *CPD*. *PRR7* is regulated by CKLs and *GR60ox2* and *CPD* are regulated by ATSKs.

Treatment of 1 μM AMI-331 significantly decreased *PRR7*, but not *GR60ox2* and *CPD*. AMI-331 at higher concentrations (5 and 25 μM) decreased *PRR7, GR60ox2*, and *CPD* (Fig. 7B). The effect of5 μM of AMI-331 on *PRR7* was stronger than those on *GR60ox2* and *CPD*. The results suggested that AMI-331 target CKLs best, but also targets ATSKs in vivo (Fig. 7C).

## Discussion

### Strong CK1 inhibitory activity of AMI-331 in vivo

We provide a novel CK1 inhibitor, namely AMI-331, that has a very strong CK1 inhibitory activity in vivo, from PHA767491 as a lead molecule. Generally, many factors such as absorption properties, transportation in and out of cells, metabolism, and inhibitory activity to targets, restrict the strength of total activity of molecular. One reason for strong in vivo CK1 inhibitory activity of AMI-331 is its in vitro CK1 inhibitory activity. However, IC_50_ of AMI-331 on CKL4 kinase activity was only 7 times stronger than PHA767491, but period-lengthening activity of AMI-331 was 100 times stronger than PHA767491. In addition, effective concentration for increasing PRR5 and TOC1 amounts by AMI-331 were about 10 to 50 μM, 10 to 50 times lower than that by PHA767491. These results suggest that other factors, such as cell membrane permeability, metabolism, and location of AMI-331 contribute to the strong AMI-331 activity in vivo.

### Selectivity of AMI-331

Our previous study suggested that PHA767491 targets the CKL family for clock regulation. However, PHA767491 can also be bound to ATSK, CPK, MPK, and other kinases. We could not estimate the selectivity of PHA767491 for CKL binding and other protein binding in a previous study (Uehara et al., 2019).

In this study, target identification of AMI-329 bead suggests that best targets of AMI-331 is the CK1 family. Adding 5 μM of AMI-331 mostly canceled binding between AMI-beads and CKL proteins (Fig. 5 and 6). Since 5 μM AMI-331 decreased binding between AMI-beads and ATSK to 50 to 20% compared to that without AMI-331, this suggests that higher concentrations of AMI-331 also binds to the ATSK family. In addition, gene expression analyses showed that 1 μM of AMI-331 affects expression of *PRR7*, which is regulated by CKLs, but not *GR60ox2* and *CPD*, which are downstream genes of ATSKs. These results suggested that AMI-331 preferentially targets CKL family rather than ATSKs. MPKs are not specifically bound by AMI-331 (Fig. 5). AT2g32850 (protein kinase) and MTK1, which were enriched by PHA767491-beads, were not enriched in AMI-beads at all. These lines of evidence suggest that the selectivity of AMI-331 toward the CKL family is enhanced compared to PHA767491.

### Possible utility for AMI-331

There are at least 14 members of the CKL family in Arabidopsis. In addition, most plants have multiple CKLs. Existence of multiple CKLs in the genome might cause difficulties in identifying physiological functions controlled by CKLs. Our previous study provided PHA767491 as a CK1 inhibitor in plants, however, the effective concentration of PHA767491 was over 40 μM. By contrast, AMI-331 effectively lengthened the clock period below 1 μM. In addition, 10 - 50 μM AMI-331 increased PRR5 and TOC1 amounts and 1 μM AMI-331 decrease *PRR7* expression. Lower effective concentration of pharmacological molecules is generally suitable for users. Using AMI-331 enables plant researchers to judge whether the CK1 family is involved in the physiological processes of interest. Note that higher concentrations of AMI-331 may modulate these physiological processes thought CKL-independent pathways. AMI-331 for basic plant research is commercially available (Tokyo Chemical Industry, product No. A3352).

## Materials and methods

### Plant materials and growth conditions

Columbia-0 (Col-0) accession plants were used as wild-type. *CCA1pro:LUC* plants reported previously (Nakamichi et al., 2005). *35Spro:PRR5-FLAG, 35Spro:PRR5-VP* and *35Spro:TOC1-VP* were reported previously (Nakamichi et al., 2016). Plants for were grown on MS containing 2% sucrose and 0.3% gellan gum under 12 h white light (70 μmol s-1 m-2) / 12 h dark conditions.

### In vitro phosphorylation assays of Arabidopsis CKL4

In vitro phosphorylation assays using recombinant CKL4 were performed as described previously (Uehara et al., 2019), with synthetic small molecules. IC261 and PF-670462 were purchased from Sigma with catalog number I0658 and SML0795, respectively. PHA767491 was synthesized as previously described (Uehara et al., 2019).

### Bioluminescence-based circadian rhythm

Bioluminescence-based circadian rhythm of *CCA1pro:LUC* treated with small molecules was analyzed by auto-luminescence detection machine (CL96, Churitsu) as described previously (Uehara et al., 2019). Period length was automatically calculated with attached software in CL96 as described (Kamioka et al., 2016).

### Synthesis of PHA767491 analogues (AMI molecules)

Synthesis of PHA767491 analogues was described in supplemental information.

### Western blotting

Four-day old seedlings grown under 12 h light/ 12 h dark (LD) were transferred into a 96-well plate by a dropper. Seedlings were treated with 20 μL of MS liquid containing 2% sucrose and AMI-331 at 2, 10, or 50 μM with a final concentration of 5% (v/v) DMSO. As a control experiment, MS containing 2% sucrose and 5% DMSO was used to treat the seedlings. Seedlings were kept under constant light (L) or constant dark (D) and frozen. Additional ‘dark’ samples of AMI-331-treated or non-treated seedlings were further kept in the dark for 24 h, and then harvested with liquid nitrogen. Frozen samples were crushed with zirconia beads (ZB-50, Tomy) in Tissue Lyser II (Qiagen). Detection of PRR5-FLAG was performed using 10-20% gradient acrylamide gel (198-15041, Wako), as previously described (Nakamichi et al., 2012). Anti-FLAG antibody (F3165, Sigma) anti-VP antibody (ab4808, Abcam) and used to detect FLAG-fusion and VP-fusion protein, respectively.

### Screening of proteins bound to AMI-329-beads

Screening of AMI-329-beads bound proteins were carried out with the method as previously described (Uehara et al., 2019). Briefly, two-week-old seedlings grown under LD conditions were harvested at different four time points (ZT2, ZT6, ZT9, and ZT17) and stored at −80 °C. Protein samples were prepared from the frozen plant samples. Prepared proteins were incubated with 0, 5, or 50 μM of AMI-331 at 4°C for 30 min with rotary mixing. Two technical replicates were used. PBS-washed AMI-329-beads were added in the protein samples, and gently rotated at 4°C for 1 h. AMI-329-beads resins were washed with bead buffer (Uehara et al., 2019) six times. The resultant resins were suspended in SDS sample buffer and boiled for 8 min. The protein sample were in-gel-digested as previously described (Uehara et al., 2019). Peptides were analyzed by a Q Exactive Hybrid Quadrupole-Orbitrap Mass Spectrometer (ThermoFisher Scientific), as described previously (Uehara et al., 2019). MS/MS spectra were interpreted and peak lists were generated using Proteome Discoverer 2.0.0.802 (ThermoFisher Scientific). Searches were performed using SEQUEST (ThermoFisher Scientific) against the *Arabidopsis thaliana* (TAIR TaxID=3702) peptide sequence database.

We validated that two technical replicates had similar results, and merged spectra data of two technical replicates. Proteins whose digested peptides’ spectra over 2 in 0 μM AMI-331 sample were selected to ensure data integrity. In the reliable group, we further selected proteins whose relative spectra (spectra in 0 μM / summed spectra in 5 and 50 μM) were over 10, as AMI-331 bound proteins (Fig. 5C). To overview spectra for potential PHA767491-target proteins (Uehara et al., 2019), spectra of these proteins were obtained from AMI-329-bead bound samples, and relative spectra against to 0 μM of AMI-331 samples were shown (Fig. 6).

### Gene expression analysis

Arabidopsis Col-0 seedlings grown under constant light conditions for X day were treated with 10 or 50 μM of Bikinin (SML0094, Sigma), 1, 5, or 20 μM of AMI-331, or DMSO as control for 6 h. Three biological replicates were used. Seedlings were frozen with liquid nitrogen and crushed with zirconia beads in a Tissue Lyser II. Powdered samples were then used for RNA isolation with illustra RNAspin (25-0500-72, GE Healthcare). RT-qPCR was performed as described previously (Nakamichi et al., 2010) using an Eco Real-Time PCR System (Illumina). Gene expression was normalized against *IPP2*, and maximal values were set to 1. Primers for detecting *IPP2, PRR7, GR60ox2*, and *CPD* are described previously (De Rybel et al., 2009; Kamioka et al., 2016).

## Accession Numbers

Sequence data for the genes described in this article is found in the Arabidopsis Information Resource under following numbers: *CCA1* (At2g46830), *CPD* (At5g05690), *IPP2* (AT3G02780) *GR60ox2* (At3g30180), and *PRR7* (At5g02810).

## Author contributions

JY and NN designed research; ANS and JY synthesized small molecules; HM, AO, and NN performed biological experiments; KK performed proteomics analysis; HM, KK, AO, TK, and NN analyzed data; JY and NN wrote the paper.

## Acknowledgements

We thank Drs. Tsuyoshi Hirota, Junya Mizoi, and Koji Takahashi for discussion of possible utilities of the AMI-331. This work was supported by Japan Society for the Promotion of Science Grants-in-Aid for Scientific Research 17K19229, 18H02136 (to NN), and 19H02726 (to JY), Grant-in-Aid for Scientific Research on Innovative Areas 15H05956 (to TK and NN), 15H05957 (to KK), 18H04428 (to JY). and Toyota Riken Scholar (to NN).

---

**Supplemental Figure 1.**
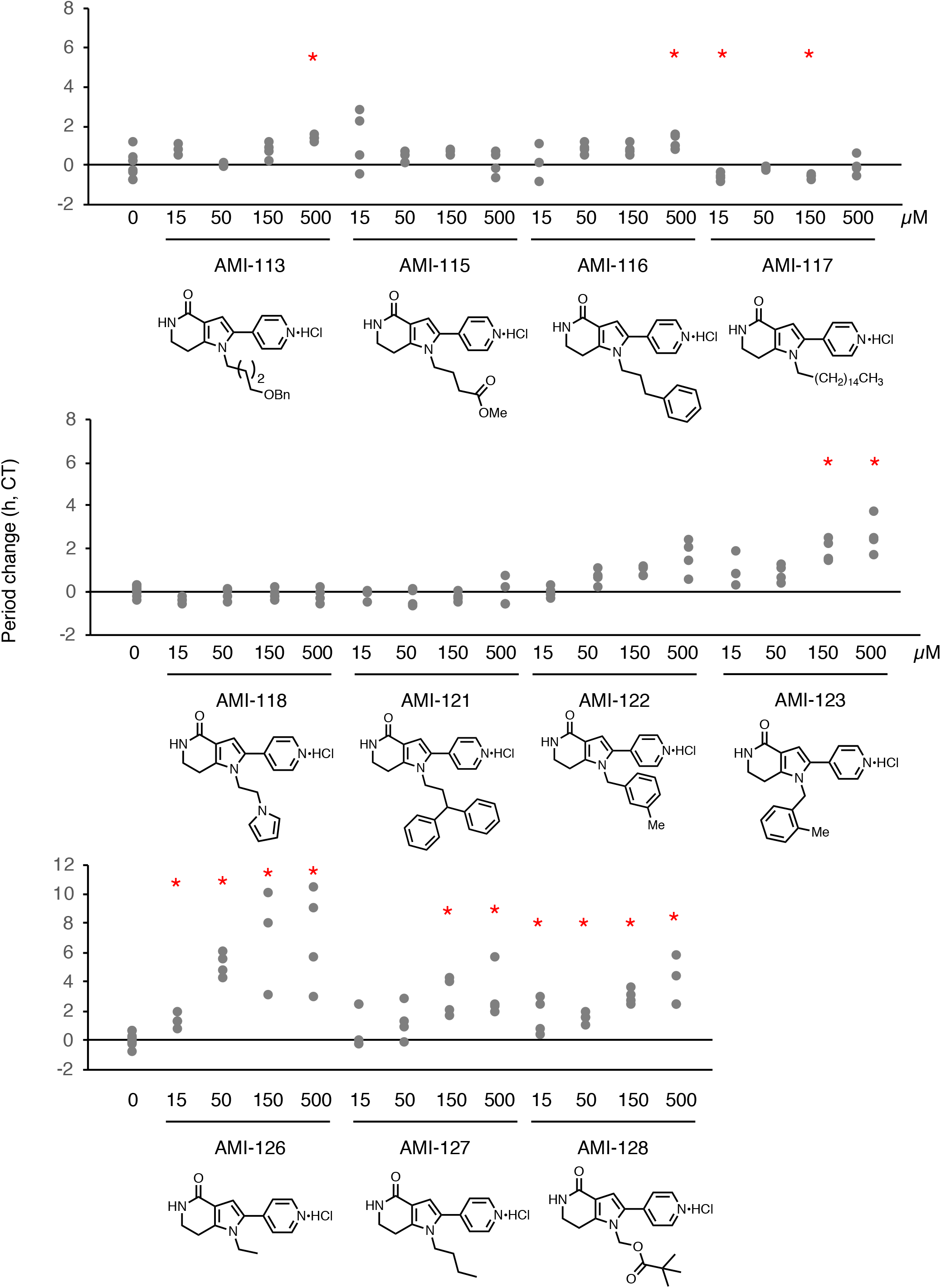
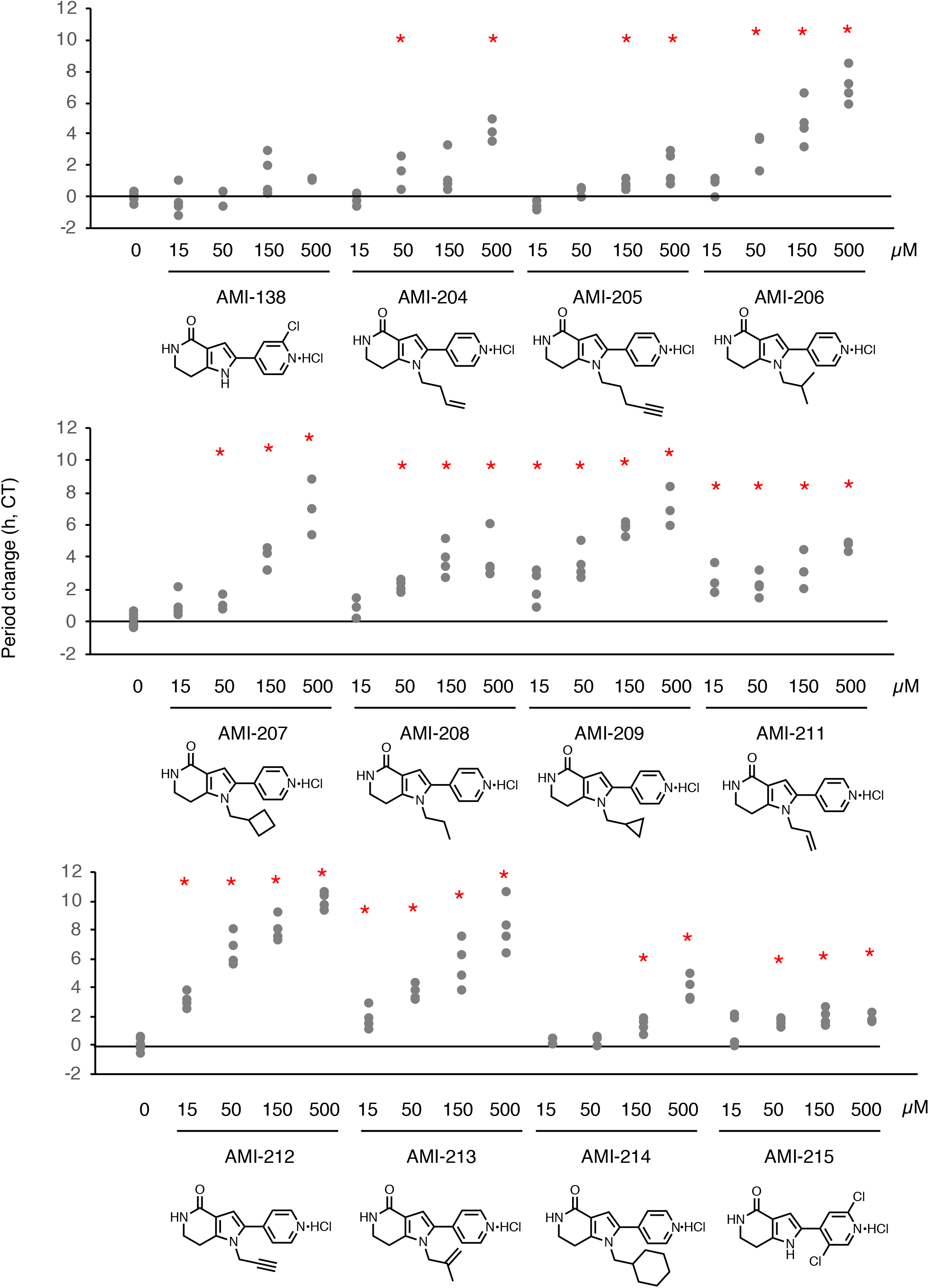
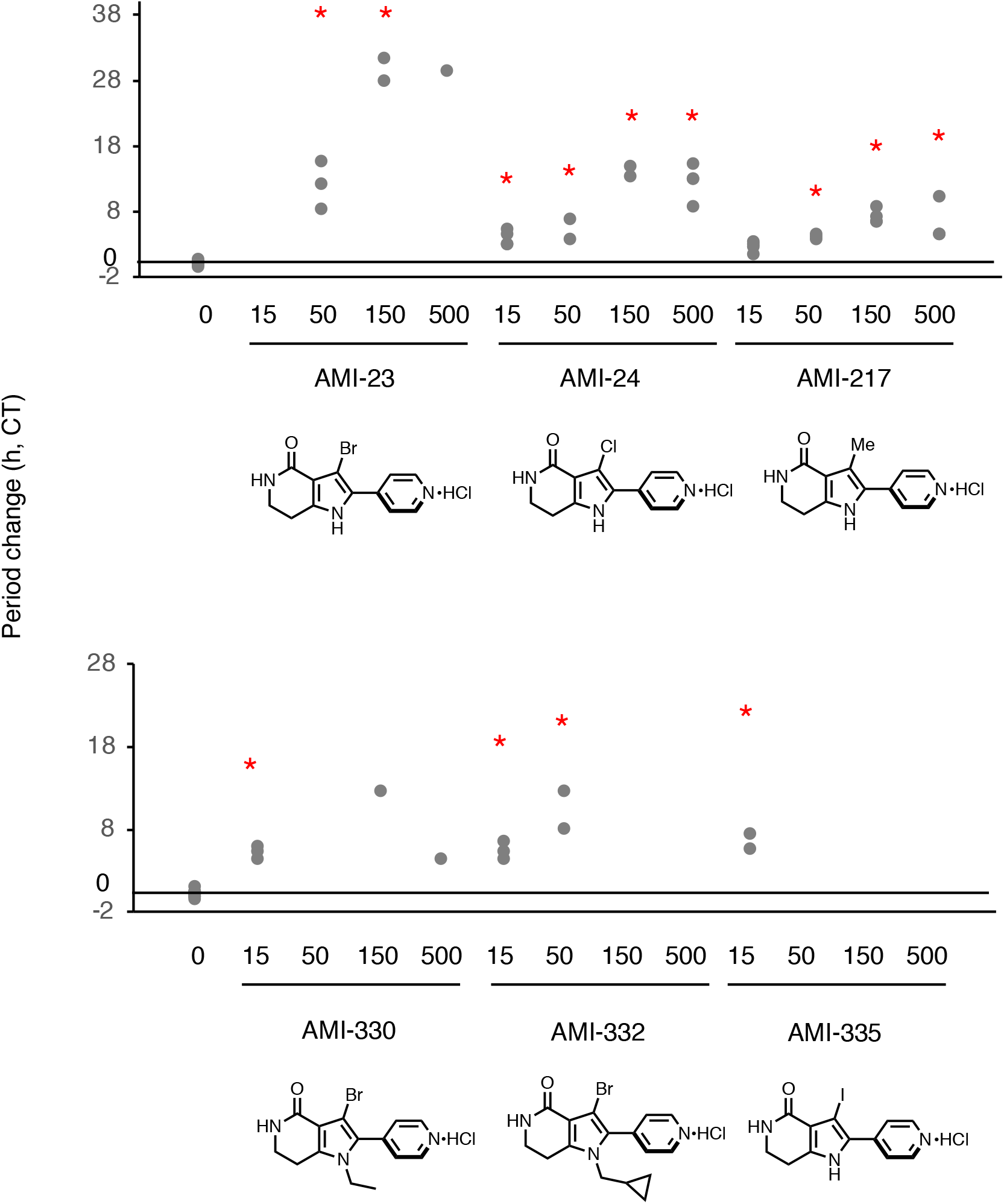
Period-lengthening activities of PHA767491-derivatives (AMI molecules). Each dot is difference of CT-corrected period length compared to DMSO control. Red asterisks show significant period change compared to DMSO samples (student’s T test *p* < 0.01).

**Supplemental Figure 2.**
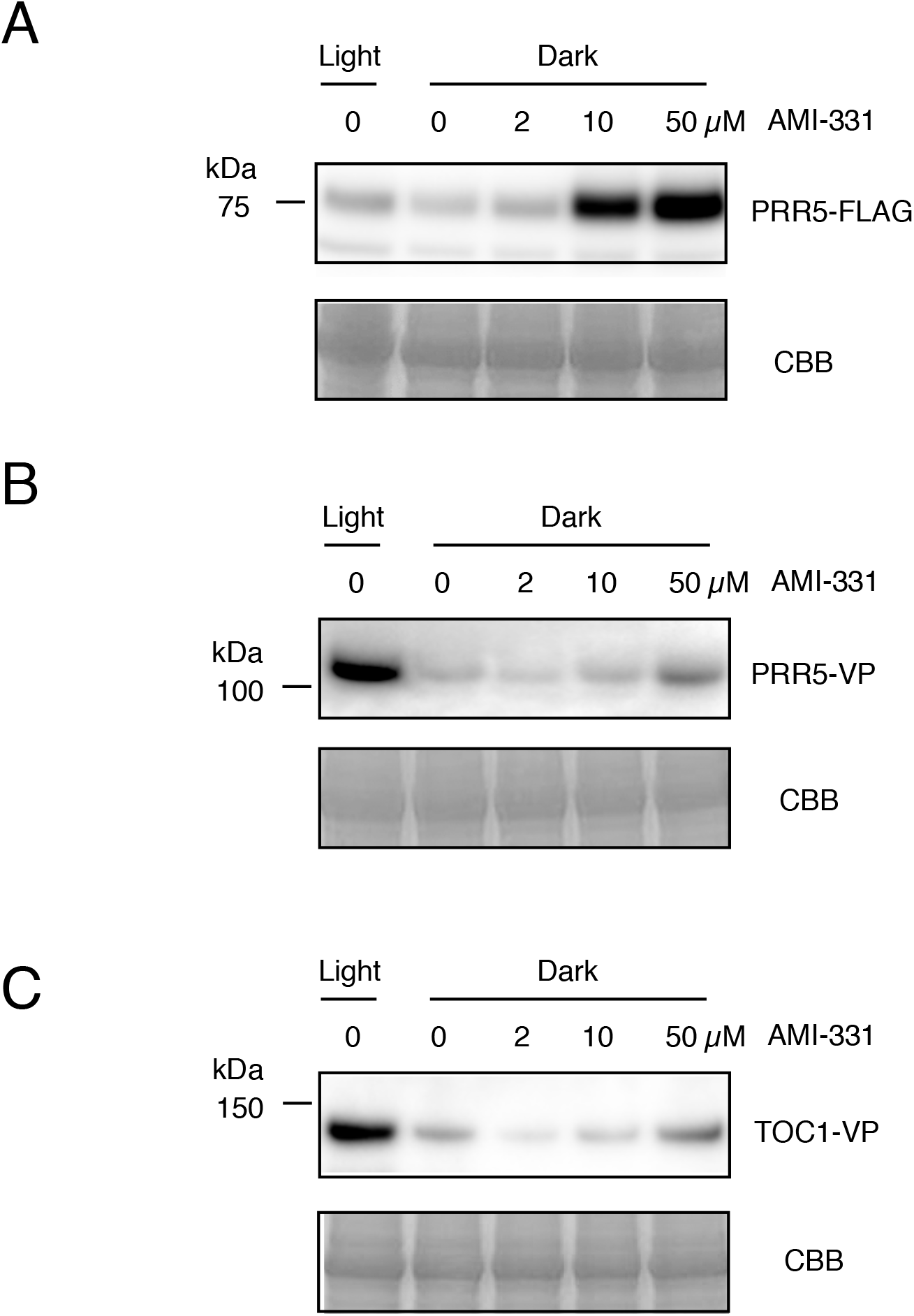
Another trial for Figure 3 (PRR5 and TOC1 proteins amount in plants treated with AMI-331). Protein amounts of PRR5-FLAG **(A)**, PRR5-VP **(B)**, and TOC1-VP **(C)** in the corresponding transgenic plants treated with AMI-331 (upper).

**Supplemental Figure 3.**
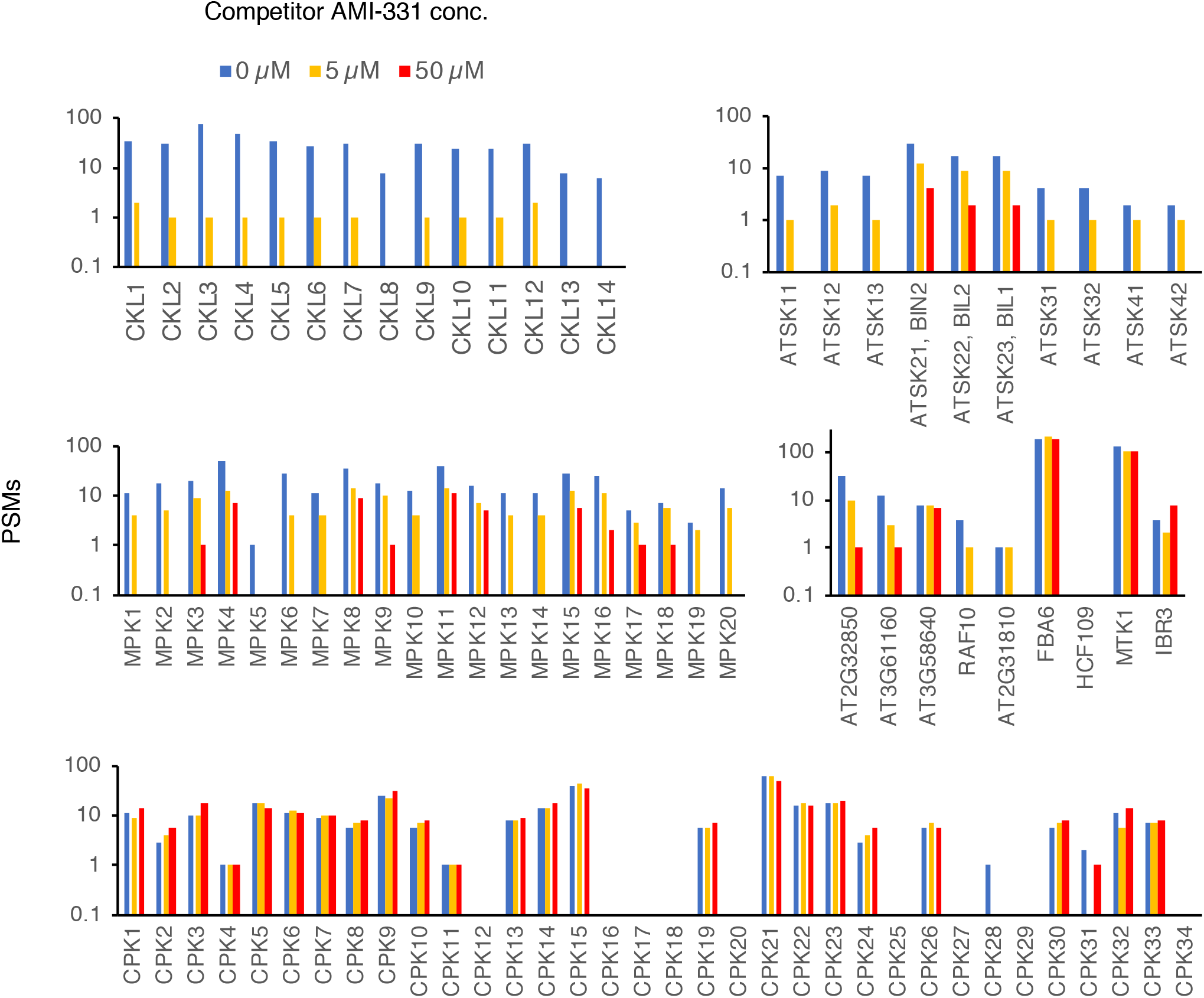
Spectra of potential PHA767491 target proteins in AMI-329-beads binding.

