## supplemental information for "Generation of strong casein kinase 1 inhibitor of *Arabidopsis thaliana*"

**Synthesis of PHA767491 derivatives and molecular probe**

1. **General**

Unless otherwise noted, all reactants or reagents including dry solvents were obtained from commercial suppliers and used as received. All work-up and purification procedures were carried out with reagent-grade solvents in air.

Analytical thin-layer chromatography (TLC) was performed using Silica gel 70 TLC Plate-Wako (0.25 mm). The developed chromatogram was analyzed by UV lamp (254 nm). Flash column chromatography was performed with Biotage Isolera^®^ equipped with Biotage SNAP Cartridge Ultra columns and Chloroform/MeOH as an eluent. Preparative thin-layer chromatography (PTLC) was performed using Wakogel B5-F silica coated plates (0.75 mm) prepared in our laboratory. High-resolution mass spectra were conducted on Thermo Fisher Scientific Exactive (ESI). Nuclear magnetic resonance (NMR) spectra were recorded on a JEOL JNM-ECZR-600 (^1^H 600 MHz, ^13^C 150 MHz). Chemical shifts for ^1^H NMR are expressed in parts per million (ppm) relative to CD_3_OD (δ 3.31 ppm). Chemical shifts for ^13^C NMR are expressed in ppm relative to CD_3_OD (49.0 ppm). Data are reported as follows: chemical shift, multiplicity (s = singlet, d = doublet, dd = doublet of doublets, ddd = doublet of doublets of doublets, t = triplet, dt = doublet of triplets, td = triplet of doublets, q = quartet, dq = doublet of quartets, m = multiplet, br s = broad singlet), coupling constant (Hz), and integration.

2. **Preparation of PHA derivatives**

**2.1 Alkylation of pyrrole at the NH position**

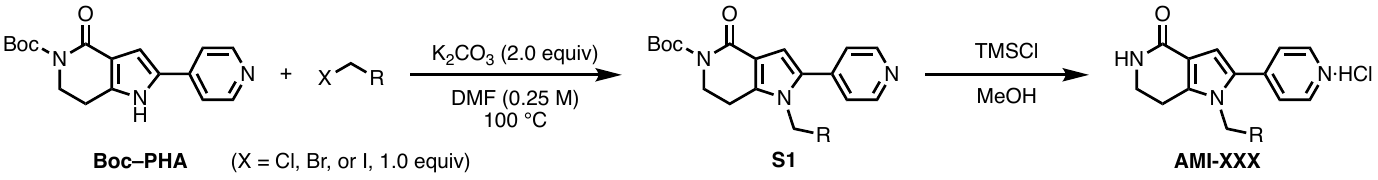

To a solution of *tert*-Butyl 4-oxo-2-(pyridin-4-yl)-1,4,6,7-tetrahydro-5*H*-pyrrolo[3,2-*c*]pyridine-5-carboxylate (**Boc–PHA**: 120 mg, 0.383 mmol, 1.0 equiv)^[1]^ and K_2_CO_3_ (105.9 mg, 0.766 mmol, 2.0 equiv) in DMF (1.5 mL) was added haloalkane (1.0 equiv) at room temperature. Then the reaction mixture was stirred at 100 °C for 24 h. It was quenched by the addition of water. The reaction mixture was extracted with ethyl acetate, washed with brine, dried over Na_2_SO_4_ and concentrated *in vacuo*. The crude mixture was purified by PTLC (CHCl_3_/MeOH = 9:1) to afford **S1**.

**S1** (10.0 mg, 1.0 equiv) in MeOH (0.3 mL) was added trimethylsilyl chloride (0.16 mL, 50.0 equiv) at room temperature. After stirring for 2 h, the reaction mixture was concentrated *in vacuo* to afford **AMI-XXX.**

**1-(4-(Benzyloxy)butyl)-2-(pyridin-4-yl)-1,5,6,7-tetrahydro-4*H*-pyrrolo[3,2-*c*]pyridin-4-one hydrochloride (AMI-113)**

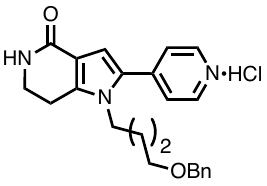

[(4-Bromobutoxy)methyl]benzene was used as the haloalkane. Purification by PTLC (CHCl_3_/MeOH = 9:1) produced **AMI-113** as a yellow solid (29% yield). ^1^H NMR (600 MHz, CD_3_OD) δ 8.58 (d, *J* = 7.2 Hz, 2H), 8.07 (d, *J* = 7.2 Hz, 2H), 7.35–7.27 (m, 5H), 7.24 (s, 1H), 4.44 (s, 2H), 4.30 (t, *J* = 7.2 Hz, 2H), 3.59 (t, *J* = 7.2 Hz, 2H), 3.48 (t, *J* = 6.0 Hz, 2H), 3.01 (t, *J* = 7.2 Hz, 2H), 1.82–1.77 (m, 2H), 1.59–1.54 (m, 2H); ^13^C NMR (150 MHz, CD_3_OD) δ 150.4, 147.0, 142.2, 139.8, 130.8, 129.5, 128.9, 128.8, 124.15, 124.10, 116.3, 73.9, 70.4, 46.7, 41.3, 28.7, 27.3, 22.7, amide carbon signal was not detected even with prolonged scans; HRMS (ESI) *m/z* = 376.2020 calcd for C_23_H_26_N_3_O_2_ [M+H]^+^, found: 376.2013.

**Methyl 4-(4-oxo-2-(pyridin-4-yl)-4,5,6,7-tetrahydro-1*H*-pyrrolo[3,2-*c*]pyridin-1-yl)butanoate hydrochloride (AMI-115)**

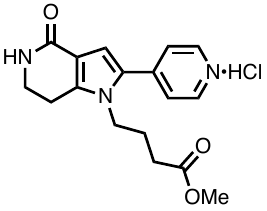

Methyl 4-chlorobutanoate was used as the haloalkane. Purification by PTLC (CHCl_3_/MeOH = 9:1) produced **AMI-115** as a yellow solid (17% yield). ^1^H NMR (600 MHz, CD_3_OD) δ 8.72 (d, *J* = 6.6 Hz, 2H), 8.17 (d, *J* = 6.6 Hz, 2H), 7.25 (s, 1H), 4.31 (t, *J* = 7.8 Hz, 2H), 3.65–3.62 (m, 4H), 3.07 (t, *J* = 7.2 Hz, 2H), 2.37 (t, *J* = 7.2 Hz, 2H), 2.00–1.94 (m, 2H); ^13^C NMR (150 MHz, CD_3_OD) δ 175.9, 150.2, 146.8, 142.43, 142.37, 130.9, 124.3, 116.1, 52.3, 46.0, 41.3, 30.9, 26.6, 22.7, amide carbon signal was not detected even with prolonged scans; HRMS (ESI) *m/z* = 314.1499 calcd for C_17_H_20_N_3_O_3_ [M+H]^+^, found: 314.1497.

**1-(3-Phenylpropyl)-2-(pyridin-4-yl)-1,5,6,7-tetrahydro-4*H*-pyrrolo[3,2-*c*]pyridin-4-one hydrochloride (AMI-116)**

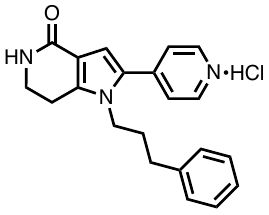

(3-Bromopropyl)benzene was used as the haloalkane. Purification by PTLC (CHCl_3_/MeOH = 9:1) produced **AMI-116** as a yellow solid (13% yield). ^1^H NMR (600 MHz, CD_3_OD) δ 8.55 (d, *J* = 7.2 Hz, 2H), 7.92 (d, *J* = 7.2 Hz, 2H), 7.26 (t, *J* = 7.2 Hz, 2H), 7.20 (s, 1H), 7.20 (t, *J* = 7.2 Hz, 1H), 7.14 (t, *J* = 7.2 Hz, 2H), 4.23 (t, *J* = 7.2 Hz, 2H), 3.60 (t, *J* = 7.2 Hz, 2H), 2.96 (t, *J* = 7.2 Hz, 2H), 2.61 (t, *J* = 7.2 Hz, 2H), 2.04–1.99 (m, 2H); ^13^C NMR (150 MHz, CD_3_OD) δ 150.3, 146.9, 142.1, 141.7, 130.8, 129.7, 129.6, 127.4, 124.1, 116.9, 116.2, 45.8, 41.2, 33.1, 32.8, 22.6, amide carbon signal was not detected even with prolonged scans ; HRMS (ESI) *m/z* = 332.1757 calcd for C_21_H_22_N_3_O [M+H]^+^, found: 332.1751.

**1-Hexadecyl-2-(pyridin-4-yl)-1,5,6,7-tetrahydro-4*H*-pyrrolo[3,2-*c*]pyridin-4-one hydrochloride (AMI-117)**

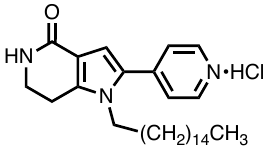

1-Chlorohexadecane was used as the haloalkane. Purification by PTLC (CHCl_3_/MeOH = 9:1) produced **AMI-117** as a yellow solid (20% yield). ^1^H NMR (600 MHz, CD_3_OD) δ 8.78 (d, *J* = 7.2 Hz, 2H), 8.15 (d, *J* = 7.2 Hz, 2H), 7.28 (s, 1H), 4.29 (t, *J* = 7.2 Hz, 2H), 3.69 (t, *J* = 7.2 Hz, 2H), 3.54 (t, *J* = 6.0 Hz, 2H), 3.10 (t, *J* = 7.2 Hz, 2H), 1.69–1.64 (m, 2H), 1.55–1.50 (m, 2H), 1.33–1.24 (m, 24H), 0.90 (t, *J* = 7.2, 1.2 Hz, 3H); ^13^C NMR (150 MHz, CD_3_OD) δ 167.6, 150.3, 147.4, 142.5, 131.4, 124.5, 116.0, 114.8, 63.0, 47.0, 41.4, 33.7, 33.1, 31.7, 30.8, 30.7, 30.63, 30.59, 30.5, 30.2, 27.5, 27.0, 23.7, 22.5, 14.5, one peak is missing due to overlapping; HRMS (ESI) *m/z* = 438.3479 calcd for C_28_H_44_N_3_O [M+H]^+^, found: 438.3473.

**1-(2-(1*H*-Pyrrol-1-yl)ethyl)-2-(pyridin-4-yl)-1,5,6,7-tetrahydro-4*H*-pyrrolo[3,2-*c*]pyridin-4-one hydrochloride (AMI-118)**

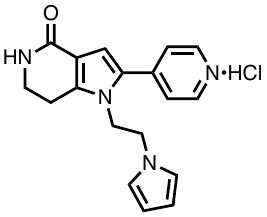

1-(2-Chloroethyl)-1*H*-pyrrole was used as the haloalkane. Purification by PTLC (CHCl_3_/MeOH = 9:1) produced **AMI-118** as a yellow solid (13% yield). ^1^H NMR (600 MHz, CD_3_OD) δ 8.64 (d, *J* = 6.0 Hz, 2H), 7.92 (d, *J* = 6.0 Hz, 2H), 7.15 (s, 1H), 6.23 (d, *J* = 3.0 Hz, 1H), 5.88 (d, *J* = 3.0 Hz, 1H), 4.58 (t, *J* = 4.8 Hz, 2H), 4.09 (t, *J* = 4.8 Hz, 2H), 3.48 (t, *J* = 7.2 Hz, 2H), 2.61 (t, *J* = 7.2 Hz, 2H); ^13^C NMR (150 MHz, CD_3_OD) δ 167.5, 149.9, 147.3, 142.0, 131.4, 124.9, 116.3, 116.0, 109.7, 109.6, 50.4, 48.4, 41.2, 22.0; HRMS (ESI) *m/z* = 307.1553 calcd for C_18_H_19_N_4_O [M+H]^+^, found: 307.1549.

**1-(3,3-Diphenylpropyl)-2-(pyridin-4-yl)-1,5,6,7-tetrahydro-4*H*-pyrrolo[3,2-*c*]pyridin-4-one hydrochloride (AMI-121)**

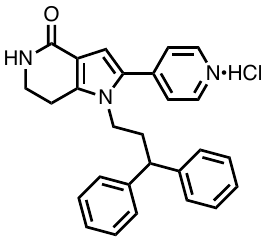

(3-Bromopropane-1,1-diyl)dibenzene was used as the haloalkane. Purification by PTLC (CHCl_3_/MeOH = 9:1) produced **AMI-121** as a yellow solid (22% yield). ^1^H NMR (600 MHz, CD_3_OD) δ 8.58 (t, *J* = 7.2 Hz, 2H), 7.93 (d, *J* = 7.2 Hz, 2H), 7.27-7.20 (m, 5H), 7.18–7.14 (m, 6H), 4.29 (t, *J* = 7.2 Hz, 2H), 3.77 (t, *J* = 7.8 Hz, 1H), 3.59 (t, *J* = 7.2 Hz, 2H), 2.86 (t, *J* = 7.2 Hz, 2H), 2.40 (q, *J* = 7.8 Hz, 2H); ^13^C NMR (150 MHz, CD_3_OD) δ 167.6, 150.1, 147.2, 144.9, 142.2, 131.4, 129.8, 128.8, 127.7, 124.6, 115.9, 115.4, 49.3, 45.5, 41.3, 36.9, 22.3; HRMS (ESI) *m/z* = 408.2070 calcd for C_27_H_26_N_3_O [M+H]^+^, found: 408.2065.

**1-(3-Methylbenzyl)-2-(pyridin-4-yl)-1,5,6,7-tetrahydro-4*H*-pyrrolo[3,2-*c*]pyridin-4-one hydrochloride (AMI-122)**

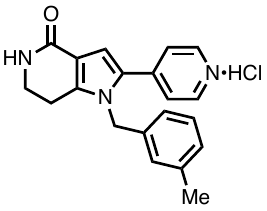

1-(Chloromethyl)-3-methylbenzene was used as the haloalkane. Purification by PTLC (CHCl_3_/MeOH = 9:1) produced **AMI-122** as a yellow solid (17% yield). ^1^H NMR (600 MHz, CD_3_OD) δ 8.48 (d, *J* = 6.6 Hz, 2H), 7.41 (d, *J* = 6.6 Hz, 1H), 7.22 (t, *J* = 7.8 Hz, 1H), 7.08 (d, *J* = 7.8 Hz, 2H), 6.87 (s, 1H), 6.81 (s, 1H), 6.72 (d, *J* = 7.8 Hz, 1H), 5.33 (s, 2H), 3.57 (t, *J* = 7.2 Hz, 2H), 2.81 (t, *J* = 7.2 Hz, 2H), 2.30 (s, 3H); ^13^C NMR (150 MHz, CD_3_OD) δ 149.8, 147.6, 142.3, 140.5, 137.4, 131.8, 130.4, 129.9, 127.4, 124.4, 123.8, 115.8, 109.1, 50.0, 41.3, 22.4, 21.4, Amide carbon signal was not detected even with prolonged scans ; HRMS (ESI) *m/z* = 318.1601 calcd for C_20_H_20_N_3_O [M+H]^+^, found: 318.1596.

**1-(2-Methylbenzyl)-2-(pyridin-4-yl)-1,5,6,7-tetrahydro-4*H*-pyrrolo[3,2-*c*]pyridin-4-one hydrochloride (AMI-123)**

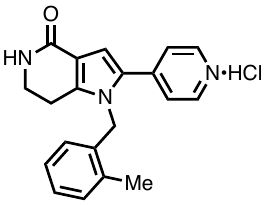

1-(Chloromethyl)-2-methylbenzene was used as the haloalkane. Purification by PTLC (CHCl_3_/MeOH = 9:1) produced **AMI-123** as a yellow solid (10% yield). ^1^H NMR (600 MHz, CD_3_OD) δ 8.60 (d, *J* = 6.6 Hz, 2H), 7.88 (d, *J* = 6.6 Hz, 2H), 7.41 (s, 1H), 7.29 (d, *J* = 7.2 Hz, 1H), 7.23 (t, *J* = 7.2 Hz, 1H), 7.16 (t, *J* = 7.2 Hz, 1H), 6.56 (d, *J* = 7.2 Hz, 1H), 5.46 (s, 2H), 3.59 (t, *J* = 7.2 Hz, 2H), 2.86 (t, *J* = 7.2 Hz, 2H), 2.37 (s, 3H); ^13^C NMR (150 MHz, CD_3_OD) δ 149.8, 142.3, 139.9, 136.4, 135.4, 132.0, 131.7, 129.2, 128.0, 125.6, 124.0, 115.8, 108.2, 41.2, 40.2, 22.4, 19.0, amide carbon signal was not detected even with prolonged scans; HRMS (ESI) *m/z* = 318.1601 calcd for C_20_H_20_N_3_O [M+H]^+^, found: 318.1596.

**1-Ethyl-2-(pyridin-4-yl)-1,5,6,7-tetrahydro-4*H*-pyrrolo[3,2-*c*]pyridin-4-one hydrochloride (AMI-126)**

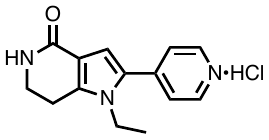

Bromoethane was used as the haloalkane. Purification by PTLC (CHCl_3_/MeOH = 9:1) produced **AMI-126** as a yellow solid (18% yield). ^1^H NMR (600 MHz, CD_3_OD) δ 8.56 (dd, *J* = 4.8, 1.2 Hz, 2H), 7.50 (dd, *J* = 4.8, 1.2 Hz, 2H), 6.72 (s, 1H), 4.12 (q, *J* = 7.2 Hz, 2H), 3.59 (t, *J* = 7.2 Hz, 2H), 2.97 (t, *J* = 7.2 Hz, 2H), 1.25 (t, *J* = 7.2 Hz, 3H); ^13^C NMR (150 MHz, CD_3_OD) δ 168.7, 150.5, 142.6, 141.9, 133.0, 124.1, 115.1, 110.0, 41.6, 40.9, 22.4, 16.4; HRMS (ESI) *m/z* = 242.1288 calcd for C_14_H_16_N_3_O [M+H]^+^, found: 242.1284.

**1-Butyl-2-(pyridin-4-yl)-1,5,6,7-tetrahydro-4*H*-pyrrolo[3,2-*c*]pyridin-4-one hydrochloride (AMI-127)**

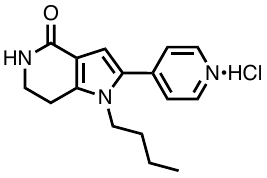

1-Iodobutane was used as the haloalkane. Purification by PTLC (CHCl_3_/MeOH = 9:1) produced **AMI-127** as a yellow solid (10% yield). ^1^H NMR (600 MHz, CD_3_OD) δ 8.58 (dd, *J* = 4.2, 1.8 Hz, 2H), 7.53 (dd, *J* = 4.2, 1.8 Hz, 2H), 6.73 (s, 1H), 4.13 (t, *J* = 7.2 Hz, 2H), 3.62 (t, *J* = 7.2 Hz, 2H), 2.99 (t, *J* = 7.2 Hz, 2H), 1.58–1.53 (m, 2H), 1.24–1.17 (m, 2H), 0.84 (t, *J* = 7.2 Hz, 3H); ^13^C NMR (150 MHz, CD_3_OD) δ 168.7, 150.5, 142.9, 142.2, 133.4, 124.2, 115.0, 110.0, 45.7, 41.6, 33.9, 22.6, 20.6, 13.8; HRMS (ESI) *m/z* = 270.1601 calcd for C_16_H_20_N_3_O [M+H]^+^, found: 270.1597.

**(4-Oxo-2-(pyridin-4-yl)-4,5,6,7-tetrahydro-1*H*-pyrrolo[3,2-*c*]pyridin-1-yl)methyl pivalate hydrochloride (AMI-128)**

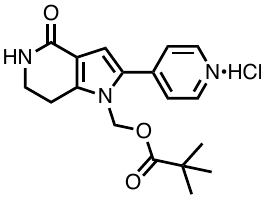

Chloromethyl pivalate was used as the haloalkane. Purification by PTLC (CHCl_3_/MeOH = 9:1) produced **AMI-128** as a yellow solid (3% yield). ^1^H NMR (600 MHz, CD_3_OD) δ 8.77 (d, *J* = 6.6 Hz, 2H), 8.15 (d, *J* = 6.6 Hz, 2H), 7.27 (s, 1H), 6.15 (s, 2H), 3.64 (t, *J* = 7.2 Hz, 2H), 3.15 (t, *J* = 7.2 Hz, 2H), 2.15 (s, 9H); ^13^C NMR (150 MHz, CD_3_OD) δ 146.5, 142.5, 142.2, 131.6, 129.0, 124.5, 124.2, 116.5, 68.2, 41.2, 39.9, 30.8, 27.2, amide carbon signal was not detected even with prolonged scans; HRMS (ESI) *m/z* = 328.1656 calcd for C_18_H_22_N_3_O_3_ [M+H]^+^, found: 328.1651.

**1-(But-3-en-1-yl)-2-(pyridin-4-yl)-1,5,6,7-tetrahydro-4*H*-pyrrolo[3,2-*c*]pyridin-4-one hydrochloride (AMI-204)**

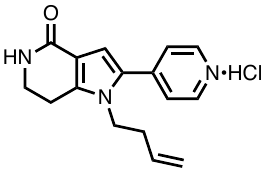

4-Bromo-1-butene was used as the haloalkane. Purification by PTLC (CHCl_3_/MeOH = 9:1) produced **AMI-204** as a yellow solid (3% yield). ^1^H NMR (600 MHz, CD_3_OD) δ 8.74 (d, *J* = 7.2 Hz, 2H), 8.14 (d, *J* = 7.2 Hz, 2H), 7.25 (s, 1H), 5.70–5.65 (m, 1H), 5.00–4.95 (m, 2H), 4.38 (t, *J* = 7.2 Hz, 2H), 3.64 (t, *J* = 7.2 Hz, 2H), 3.08 (t, *J* = 7.2 Hz, 2H), 2.40 (q, *J* = 7.2 Hz, 2H); ^13^C NMR (150 MHz, CD_3_OD) δ 167.6, 150.5, 147.4, 142.5, 134.6, 131.3, 124.5, 119.1, 116.1, 115.3, 46.3, 41.4, 35.8, 22.7; HRMS (ESI) *m/z* = 268.1444 calcd for C_16_H_18_N_3_O [M+H]^+^, found: 268.1440.

**1-(Pent-4-yn-1-yl)-2-(pyridin-4-yl)-1,5,6,7-tetrahydro-4*H*-pyrrolo[3,2-*c*]pyridin-4-one hydrochloride (AMI-205)**

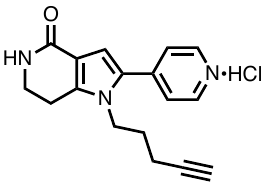

5-Chloro-1-pentyne was used as the haloalkane. Purification by PTLC (CHCl_3_/MeOH = 9:1) produced **AMI-205** as a yellow solid (6% yield). ^1^H NMR (600 MHz, CD_3_OD) δ 8.75 (d, *J* = 7.2 Hz, 2H), 8.17 (d, *J* = 7.2 Hz, 2H), 7.27 (s, 1H), 4.43 (t, *J* = 7.2 Hz, 2H), 3.68 (t, *J* = 7.2 Hz, 2H), 3.14–3.10 (m, 2H), 2.35 (t, *J* = 3.0 Hz, 1H), 2.16 (td, *J* = 6.0, 3.0 Hz, 2H), 1.90–1.85 (m, 2H); ^13^C NMR (150 MHz, CD_3_OD) δ 166.3, 150.3, 147.4, 142.4, 131.3, 126.7, 124.7, 116.1, 83.2, 71.3, 45.6, 41.4, 30.2, 22.6, 15.9; HRMS (ESI) *m/z* = 280.1444 calcd for C_17_H_18_N_3_O [M+H]^+^, found: 280.1440.

**1-Isopentyl-2-(pyridin-4-yl)-1,5,6,7-tetrahydro-4*H*-pyrrolo[3,2-*c*]pyridin-4-one hydrochloride (AMI-206)**

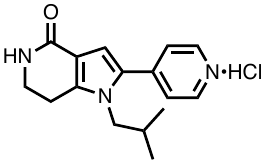

1-Bromo-2-methylpropane was used as the haloalkane. Purification by PTLC (CHCl_3_/MeOH = 9:1) produced **AMI-206** as a yellow solid (11% yield). ^1^H NMR (600 MHz, CD_3_OD) δ 8.56 (d, *J* = 6.0 Hz, 2H), 7.52 (d, *J* = 6.0 Hz, 2H), 6.72 (s, 1H), 3.98 (d, *J* = 7.8 Hz, 2H), 3.59 (t, *J* = 7.2 Hz, 2H), 2.95 (t, *J* = 7.2 Hz, 2H), 1.71–1.64 (m, 1H), 0.71 (d, *J* = 7.2 Hz, 6H); ^13^C NMR (150 MHz, CD_3_OD) δ 168.7, 150.5, 143.3, 142.6, 133.6, 124.2, 114.9, 110.4, 53.1, 41.7, 31.1, 23.0, 19.8; HRMS (ESI) *m/z* = 270.1601 calcd for C_16_H_20_N_3_O [M+H]^+^, found: 270.1598.

**1-Isobutyl-2-(pyridin-4-yl)-1,5,6,7-tetrahydro-4*H*-pyrrolo[3,2-*c*]pyridin-4-one hydrochloride (AMI-207)**

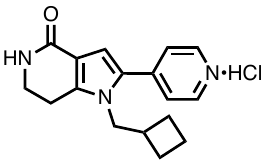

(Bromomethyl)cyclobutane was used as the haloalkane. Purification by PTLC (CHCl_3_/MeOH = 9:1) produced **AMI-207** as a yellow solid (14% yield). ^1^H NMR (600 MHz, CD_3_OD) δ 8.59 (dd, *J* = 4.8, 2.4 Hz, 2H), 7.53 (dd, *J* = 4.8, 2.4 Hz, 2H), 6.72 (s, 1H), 4.20 (d, *J* = 6.6 Hz, 2H), 3.61 (t, *J* = 6.6 Hz, 2H), 2.99 (t, *J* = 7.2 Hz, 2H), 2.48–2.40 (m, 1H), 1.87–1.81 (m, 2H), 1.81–1.74 (m, 1H), 1.73–1.67 (m, 1H), 1.57–1.51 (m, 2H); ^13^C NMR (150 MHz, CD_3_OD) δ 168.7, 150.4, 143.2, 142.3, 133.4, 124.6, 115.0, 110.3, 50.7, 41.7, 38.1, 27.0, 23.0, 18.9; This compound was not ionized by ESI and DART mass.

**1-Propyl-2-(pyridin-4-yl)-1,5,6,7-tetrahydro-4*H*-pyrrolo[3,2-*c*]pyridin-4-one hydrochloride (AMI-208)^1^**

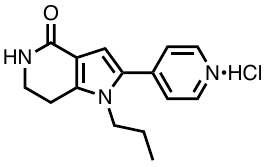

1-Chloropropane was used as the haloalkane. Purification by PTLC (CHCl_3_/MeOH = 9:1) produced **AMI-208** as a yellow solid (1% yield). ^1^H NMR (600 MHz, CD_3_OD) δ 8.66 (brs, 2H), 7.28 (d, *J* = 6.0 Hz, 2H), 6.78 (s, 1H), 4.14 (t, *J* = 6.0 Hz, 2H), 3.92 (t, *J* = 7.2 Hz, 2H), 2.91 (t, *J* = 6.0 Hz, 2H), 1.60–1.53 (m, 11H), 0.79 (t, *J* = 7.2 Hz, 3H); ^13^C NMR (150 MHz, CD_3_OD) δ 162.3, 153.6, 150.1, 140.5, 140.0, 133.0, 122.7, 115.6, 109.9, 82.5, 46.4, 44.7, 28.1, 24.1, 22.4, 10.8; HRMS (DART) *m/z* = 356.1974 calcd for C_15_H_18_N_3_O [M+H]^+^, found: 356.1976.

**1-(Cyclopropylmethyl)-2-(pyridin-4-yl)-1,5,6,7-tetrahydro-4*H*-pyrrolo[3,2-*c*]pyridin-4-one hydrochloride (AMI-209)**

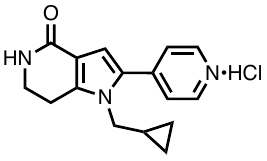

(Chloromethyl)cyclopropane was used as the haloalkane. Purification by PTLC (CHCl_3_/MeOH = 9:1) produced **AMI-209** as a yellow solid (7% yield). ^1^H NMR (600 MHz, CD_3_OD) δ 8.58 (dd, *J* = 4.8, 1.8 Hz, 2H), 8.18–8.22 (dd, *J* = 4.8, 1.8 Hz, 2H), 6.74 (s, 1H), 4.04 (d, *J* = 6.6 Hz, 2H), 3.62 (t, *J* = 7.2 Hz, 2H), 3.01(t, *J* = 7.2 Hz, 2H), 1.00–0.91 (m, 1H), 0.49–0.44 (m, 2H), 0.11–0.07 (m, 2H); ^13^C NMR (150 MHz, CD_3_OD) δ 168.7, 150.5, 143.1, 142.1, 133.4, 124.5, 115.1, 110.1, 49.9, 41.7, 22.9, 12.9, 4.3; HRMS (ESI) *m/z* = 268.1444 calcd for C_16_H_18_N_3_O [M+H]^+^, found: 268.1444.

**1-Allyl-2-(pyridin-4-yl)-1,5,6,7-tetrahydro-4*H*-pyrrolo[3,2-*c*]pyridin-4-one hydrochloride (AMI-211)**

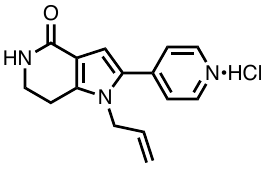

Allyl bromide was used as the haloalkane. Purification by PTLC (CHCl_3_/MeOH = 9:1) produced **AMI-211** as a yellow solid (3% yield). ^1^H NMR (600 MHz, CD_3_OD) δ 8.69 (d, *J* = 7.2 Hz, 2H), 8.07 (d, *J* = 7.2 Hz, 2H), 7.33 (s, 1H), 6.18–6.13 (m, 1H), 5.36 (d, *J* = 10.8 Hz, 1H), 4.93–4.84 (m, 3H), 3.62 (t, *J* = 6.6 Hz, 2H), 2.97 (t, *J* = 6.6 Hz, 2H); ^13^C NMR (150 MHz, CD_3_OD) δ 149.9, 147.1, 142.3, 134.5, 131.2, 124.1, 117.5, 116.9, 115.7, 111.9, 41.2, 22.3, amide carbon signal was not detected even with prolonged scans; HRMS (ESI) *m/z* = 254.1288 calcd for C_15_H_16_N_3_O [M+H]^+^, found: 254.1284.

**1-(Prop-2-yn-1-yl)-2-(pyridin-4-yl)-1,5,6,7-tetrahydro-4*H*-pyrrolo[3,2-*c*]pyridin-4-one hydrochloride (AMI-212)**

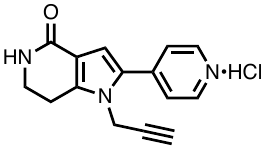

Propargyl chloride was used as the haloalkane. Purification by PTLC (CHCl_3_/MeOH = 9:1) produced **AMI-212** as a yellow solid (11% yield). ^1^H NMR (600 MHz, CD_3_OD) δ 8.61 (dd, *J* = 4.8, 1.2 Hz, 2H), 7.62 (dd, *J* = 4.8, 1.2 Hz, 2H), 6.79 (s, 1H), 4.86 (d, *J* = 2.4 Hz, 2H), 3.64 (t, *J* = 7.2 Hz, 2H), 3.07–3.04 (m, 3H); ^13^C NMR (150 MHz, CD_3_OD) δ 168.4, 150.6, 142.4, 141.8, 133.4, 123.9, 115.6, 109.8, 78.9, 75.9, 41.4, 35.7, 22.3; HRMS (ESI) *m/z* = 252.1131 calcd for C_15_H_14_N_3_O [M+H]^+^, found: 252.1131.

**1-(2-methylallyl)-2-(pyridin-4-yl)-1,5,6,7-tetrahydro-4*H*-pyrrolo[3,2-*c*]pyridin-4-one hydrochloride (AMI-213)**

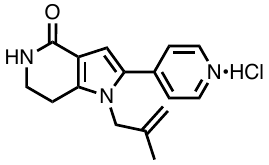

3-Chloro-2-methyl-1-propene was used as the haloalkane. Purification by PTLC (CHCl_3_/MeOH = 9:1) produced **AMI-213** as a yellow solid (7% yield). ^1^H NMR (600 MHz, CD_3_OD) δ 8.55 (dd, *J* = 4.8, 2.4 Hz, 2H), 7.49 (dd, *J* = 4.8, 2.4 Hz, 2H), 6.82 (s, 1H), 4.99 (s, 1H), 4.59 (s, 2H), 4.40 (s, 1H), 3.60 (t, *J* = 7.2 Hz, 2H), 2.88 (t, *J* = 7.2 Hz, 2H), 1.77 (s, 3H); ^13^C NMR (150 MHz, CD_3_OD) δ 149.9, 147.2, 142.6, 142.3, 131.3, 126.7, 124.0, 116.6, 115.6, 112.3, 51.8, 41.3, 22.3, 20.0, amide carbon signal was not detected even with prolonged scans; HRMS (ESI) *m/z* = 268.1444 calcd for C_16_H_18_N_3_O [M+H]^+^, found: 268.1440.

**1-(Cyclohexylmethyl)-2-(pyridin-4-yl)-1,5,6,7-tetrahydro-4*H*-pyrrolo[3,2-*c*]pyridin-4-one hydrochloride (AMI-214)**

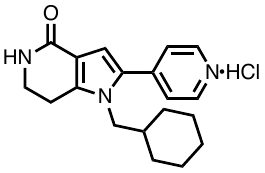

(Bromomethyl)cyclohexane was used as the haloalkane. Purification by PTLC (CHCl_3_/MeOH = 9:1) produced **AMI-214** as a yellow solid (24% yield). ^1^H NMR (600 MHz, CD_3_OD) δ 8.72 (d, *J* = 7.2 Hz, 2H), 8.15 (d, *J* = 7.2 Hz, 2H), 7.24 (s, 1H), 4.16 (d, *J* = 7.8 Hz, 2H), 3.62 (t, *J* = 7.2 Hz, 2H), 3.04 (t, *J* = 7.2 Hz, 2H), 1.64–1.58 (m, 3H), 1.46–1.39 (m, 3H), 1.12–1.04 (m, 3H), 0.87 (q, *J* = 11.4 Hz, 2H); ^13^C NMR (150 MHz, CD_3_OD) δ 167.6, 150.9, 147.6, 142.4, 131.3, 124.3, 116.5, 116.0, 52.8, 41.4, 40.7, 31.3, 27.1, 26.6, 23.1; HRMS (ESI) *m/z* = 310.1914 calcd for C_19_H_24_N_3_O [M+H]^+^, found: 310.1909.

**2.2. Substitution of pyrrole at C3-position**

**3-Bromo-2-(pyridin-4-yl)-1,5,6,7-tetrahydro-4*H*-pyrrolo[3,2-*c*]pyridin-4-one hydrochloride (AMI-23)**

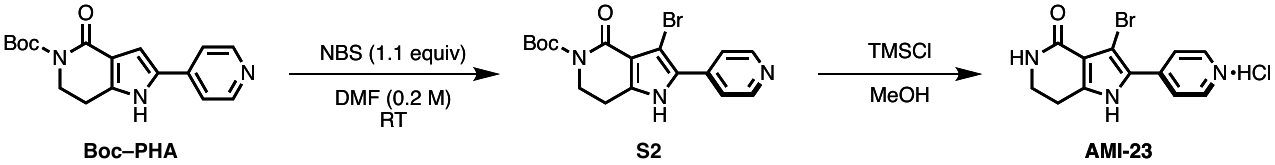

To a solution of **Boc–PHA** (20 mg, 63.8 μmol, 1.0 equiv) in DMF (0.3 mL) was added *N*-bromosuccinimide (NBS: 12.5 mg, 70.2 μmol, 1.1 equiv) at room temperature. It was quenched by the addition of Hünig's base. The reaction mixture was extracted with ethyl acetate, washed with brine, dried over Na_2_SO_4_ and concentrated *in vacuo.* The crude mixture was purified by PTLC (CHCl_3_/MeOH = 9:1) to afford **S2** as a yellow oil (13.9 mg, 54% yield).

**S2** (10.0 mg, 1.0 equiv) in MeOH (0.3 mL) was added trimethylsilyl chloride (0.16 mL, 50 equiv) at room temperature. After stirring for 2 h, the reaction mixture was concentrated *in vacuo* to afford **AMI-23** as a yellow solid. ^1^H NMR (600 MHz, D_2_O) δ 8.53 (d, *J* = 7.2 Hz, 2H), 8.19 (d, *J* = 7.2 Hz, 2H), 3.47 (t, *J* = 7.2 Hz, 2H), 2.89 (t, *J* = 7.2 Hz, 2H); ^13^C NMR (150 MHz, D_2_O) δ 165.6, 146.3, 143.6, 140.4, 123.0, 120.4, 113.5, 111.7, 39.3, 21.3; HRMS (ESI) *m/z* = 292.0080 calcd for C_12_H_11_BrN_3_O [M+H]^+^, found: 292.0080.

**3-Chloro-2-(pyridin-4-yl)-1,5,6,7-tetrahydro-4*H*-pyrrolo[3,2-*c*]pyridin-4-one hydrochloride (AMI-24)**

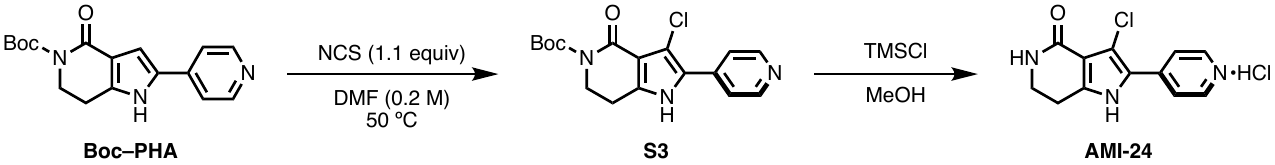

To a solution of **Boc–PHA** (20 mg, 63.8 μmol, 1.0 equiv) in DMF (0.3 mL) was added *N*-chlorosuccinimide (NCS: 9.4 mg, 70.2 μmol, 1.1 equiv) at 50 °C. It was quenched by the addition of Hünig's base. The reaction mixture was extracted with ethyl acetate, washed with brine, dried over Na_2_SO_4_ and concentrated *in vacuo.* The crude mixture was purified by PTLC (CHCl_3_/MeOH = 9:1) to afford **S3** as a yellow oil (11 mg, 50% yield).

**S3** (10.0 mg, 1.0 equiv) in MeOH (0.3 mL) was added trimethylsilyl chloride (0.16 mL, 50 equiv) at room temperature. After stirring for 2 h, the reaction mixture was concentrated *in vacuo* to afford **AMI-24** as a yellow solid. ^1^H NMR (600 MHz, D_2_O) δ 8.54 (dd, *J* = 7.2, 2.4 Hz, 2H), 8.16 (dd, *J* = 7.2, 2.4 Hz, 2H), 3.47 (td, *J* = 7.2, 1.8 Hz, 2H), 2.91 (td, *J* = 7.2,1.8 Hz, 2H); ^13^C NMR (150 MHz, D_2_O) δ 165.6, 145.9, 143.1, 140.4, 121.7, 120.1, 116.9, 112.0, 39.5, 21.3; HRMS (ESI) *m/z* = 248.0585 calcd for C_12_H_11_ClN_3_O [M+H]^+^, found: 248.0586.

**3-Methyl-2-(pyridin-4-yl)-1,5,6,7-tetrahydro-4*H*-pyrrolo[3,2-*c*]pyridin-4-one hydrochloride (AMI-217)**

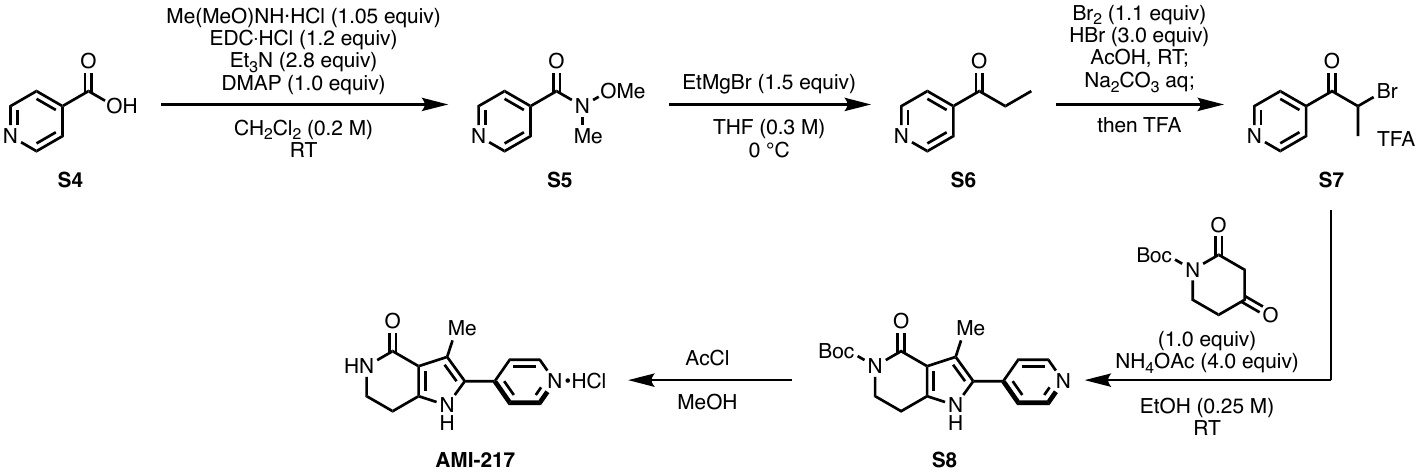

To a round bottomed flask with 4-pyridinecarboxylic acid (**S4**: 500 mg, 4.06 mmol, 1.0 equiv) were added EDC·HCl (934 mg, 4.87 mmol, 1.2 equiv), *N,N*-dimethyl-4-aminopyridine (DMAP: 496mg, 4.06 mmol, 1.0 equiv), Me(MeO)NH·HCl (416 mg, 4.26 mmol, 1.05 equiv), Et_3_N (1.15 g, 11.4 mmol, 2.8 equiv), and CH_2_Cl_2_ (20 mL). After stirring the mixture at room temperature for several hours with monitoring reaction progress with TLC, the reaction was quenched with sat. NaHCO_3_ aq. and extracted three times with CH_2_Cl_2_, and washed with brine. The combined organic layer was dried over Na_2_SO_4_, filtrated, and concentrated *in vacuo*. The residue was purified by flash column chromatography to afford amide **S5**.

A solution of EtMgBr (1 M in THF, 5.42 mL, 5.42 mmol, 1.5 equiv) was added slowly to a stirred solution of **S5** (600 mg, 3.61 mmol, 1.0 equiv) in dry THF (12 mL) at 0 °C, and the mixture was stirred at 0 °C. The reaction mixture was quenched with sat. NH_4_Cl aq. and the mixture extracted with EtOAc. The combined organic fraction was washed with water, washed with brine, and dried over Na_2_SO_4_, and the solvent was concentrated *in vacuo*. The residue was purified by column chromatography to afford ketone **S6**.

Br_2_ (473 mg, 2.96 mmol, 1.1 equiv) was added dropwise to a stirred solution of **S6** (364 mg, 2.96 mmol, 1.0 equiv) in 5.1 M HBr/HOAc (1.56 mL) at room temperature. The reaction mixture was quenched with sat. Na_2_CO_3_ aq. and extracted three times with CH_2_Cl_2_. The combined organic layer was dried over Na_2_SO_4_, added TFA, filtrated, and concentrated *in vacuo* to give **S7** as the TFA salt.

To a solution of 2,4-dioxopiperidine-1-carboxylate (575 mg, 2.69 mmol, 1.0 equiv) and **S7** (884 mg, 2.69 mmol, 1.0 equiv) in EtOH (10.8 mL) was added ammonium acetate (831 mg, 10.8 mmol, 4.0 equiv) at room temperature. After stirring, the reaction mixture was quenched with sat. NaHCO_3_ aq., then the layers were separated. The aqueous layer was extracted with ethyl acetate and the combined organic layer was dried over Na_2_SO_4_ and concentrated *in vacuo*. The residue was purified by column chromatography to afford **S8** (12% yield).

**S8** (10.0 mg, 1.0 equiv) in MeOH (0.3 mL) was added trimethylsilyl chloride (0.16 mL, 50 equiv) at room temperature. After stirring for 2 h, the reaction mixture was concentrated *in vacuo* to afford **AMI-217** as a yellow solid. ^1^H NMR (600 MHz, CD_3_OD) δ 8.48 (dd, *J* = 4.8, 1.8 Hz, 2H), 7.51 (dd, *J* = 4.8, 1.8 Hz, 2H), 3.54 (t, *J* = 7.2 Hz, 2H), 2.91 (t, *J* = 7.2 Hz, 2H), 2.53 (s, 3H); ^13^C NMR (150 MHz, CD_3_OD) 169.6, 150.2, 142.7, 140.3, 126.6, 121.6, 121.3, 114.6, 41.6, 23.1, 11.7. HRMS (ESI) *m/z* = 228.1131 calcd for C_13_H_14_N_3_O [M+H]^+^, found: 228.1130.

**2.3 Synthesis of hybrid derivatives**

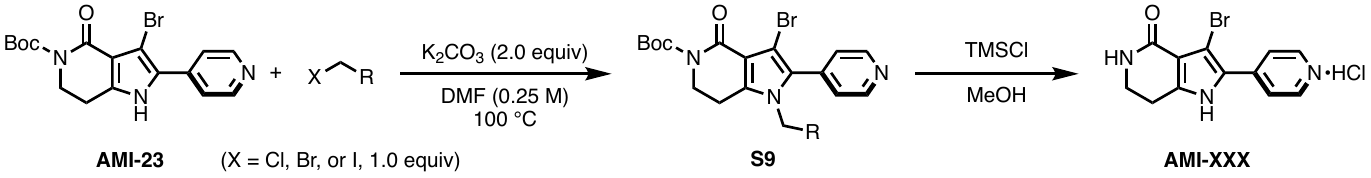

To a solution of *tert*-Butyl 3-bromo-2-(pyridin-4-yl)-1,5,6,7-tetrahydro-4*H*-pyrrolo[3,2-*c*]pyridin-4-one (**AMI-23**: 120 mg, 0.383 mmol, 1.0 equiv) and K_2_CO_3_ (105.9 mg, 0.766 mmol, 2.0 equiv) in DMF (1.5 mL) was added haloalkane (1.0 equiv) at room temperature. Then the reaction mixture was stirred at 100 °C for 24 h. It was quenched by the addition of water. The reaction mixture was extracted with ethyl acetate, washed with brine, dried over Na_2_SO_4_ and concentrated *in vacuo*. The crude mixture was purified by PTLC (CHCl_3_/MeOH = 9:1) to afford **S9**.

**S9** (10.0 mg, 1.0 equiv) in MeOH (0.3 mL) was added trimethylsilyl chloride (0.16 mL, 50.0 equiv) at room temperature. After stirring for 2 h, the reaction mixture was concentrated *in vacuo* to afford **AMI-330**, **AMI-331**, and **AMI-332.**

**3-Bromo-1-ethyl-2-(pyridin-4-yl)-1,5,6,7-tetrahydro-4*H*-pyrrolo[3,2-*c*]pyridin-4-one hydrochloride (AMI-330)**

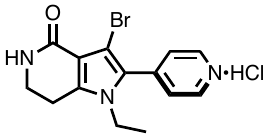

Bromoethane was used as the haloalkane. Purification by PTLC (CHCl_3_/MeOH = 9:1) produced **AMI-330** as a yellow solid (55% yield). ^1^H NMR (600 MHz, CD_3_OD) δ 8.91 (d, *J* = 7.2 Hz, 2H), 8.21 (d, *J* = 7.2 Hz, 2H), 4.16 (q, *J* = 7.2 Hz, 2H), 3.60 (t, *J* = 7.2 Hz, 2H), 3.06 (t, *J* = 7.2 Hz, 2H), 1.22 (t, *J* = 7.2 Hz, 3H); ^13^C NMR (150 MHz, CD_3_OD) δ 166.2, 149.4, 144.0, 142.8, 128.6, 128.2, 114.4, 101.2, 42.1, 40.8, 22.7, 16.2; HRMS (ESI) *m/z* = 320.0393 calcd for C_14_H_15_BrN_3_O [M+H]^+^, found: 320.0391.

**3-Bromo-1-(prop-2-yn-1-yl)-2-(pyridin-4-yl)-1,5,6,7-tetrahydro-4*H*-pyrrolo[3,2-*c*]pyridin-4-one hydrochloride (AMI-331)**

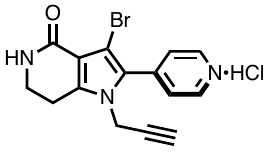

Propargyl chloride was used as the haloalkane. Purification by PTLC (CHCl_3_/MeOH = 9:1) produced **AMI-331** as a yellow solid (44% yield). ^1^H NMR (600 MHz, CD_3_OD) δ 8.68 (dd, *J* = 7.2, 3.0 Hz, 2H), 7.59 (dd, *J* = 7.2, 3.0 Hz, 2H), 4.72 (d, *J* = 3.6 Hz, 2H), 3.59 (t, *J* = 10.2 Hz, 2H), 3.04 (t, *J* = 10.2 Hz, 2H), 2.98 (t, *J* = 3.6 Hz, 1H); ^13^C NMR (150 MHz, CD_3_OD) δ 168.4, 150.6, 142.4, 141.8, 133.4, 123.9, 115.6, 109.8, 78.9, 75.9, 41.4, 35.7, 22.3; HRMS (ESI) *m/z* = 330.0237 calcd for C_15_H_13_BrN_3_O [M+H]^+^, found: 330.0233.

**3-Bromo-1-(cyclopropylmethyl)-2-(pyridin-4-yl)-1,5,6,7-tetrahydro-4*H*-pyrrolo[3,2-*c*]pyridin-4-one hydrochloride (AMI-332)**

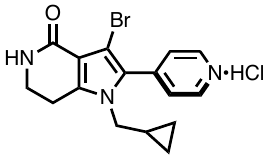

(Chloromethyl)cyclopropane was used as the haloalkane. Purification by PTLC (CHCl_3_/MeOH = 9:1) produced **AMI-332** as a yellow solid (24% yield). ^1^H NMR (600 MHz, CD_3_OD) δ 8.88 (d, *J* = 6.0 Hz, 2H), 8.22 (d, *J* = 6.0 Hz, 2H), 4.01 (d, *J* = 6.6 Hz, 2H), 3.60 (t, *J* = 6.6 Hz, 2H), 3.08 (t, *J* = 6.6 Hz, 2H), 0.92–0.85 (m, 1H), 0.45 (d, *J* = 7.2 Hz, 2H), 0.09 (t, *J* = 3.6 Hz, 2H); ^13^C NMR (150 MHz, CD_3_OD) δ 166.2, 150.0, 144.5, 142.8, 128.8, 128.6, 114.3, 101.5, 51.3, 40.9, 23.2, 12.9, 4.6; HRMS (ESI) *m/z* = 346.0550 calcd for C_16_H_17_BrN_3_O [M+H]^+^, found: 346.0549.

**2.4. Conversion of the pyridine^[^**^2]^

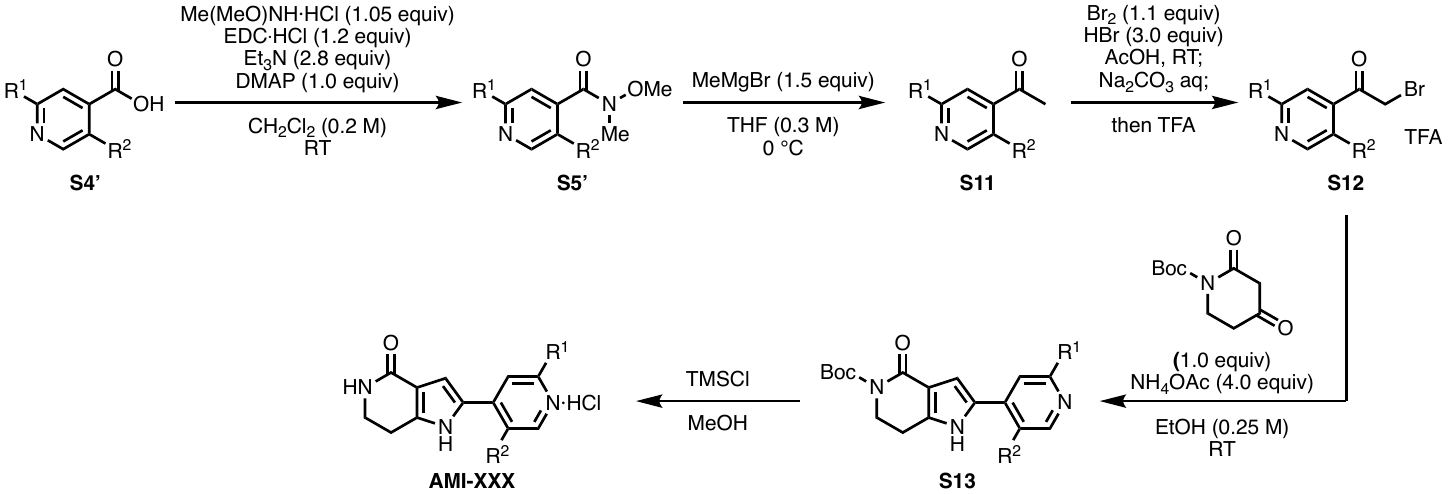

To a round bottomed flask with 4-pyridinecarboxylic acid (**S4’**: 200 mg, 1.0 equiv) were added EDC·HCl (1.2 equiv), DMAP (1.0 equiv), Me(MeO)NH·HCl (1.05 equiv), Et_3_N (2.8 equiv), and CH_2_Cl_2_ (0.2 M). After stirring the mixture at room temperature for several hours with monitoring reaction progress with TLC, the reaction was quenched with sat. NaHCO_3_ aq. and extracted three times with CH_2_Cl_2_, and washed with brine. The combined organic layer was dried over Na_2_SO_4_, filtrated, and concentrated *in vacuo*. The residue was purified by flash column chromatography to afford amide **S5’**.

A solution of MeMgBr (1 M in THF, 1.5 equiv) was added slowly to a stirred solution of **S5’** (1.0equiv) in dry THF (0.3 M) at 0 °C, and the mixture was stirred at 0 °C. The reaction mixture was quenched with sat. NH_4_Cl aq. and the mixture extracted with EtOAc. The combined organic fraction was washed with water, washed with brine, and dried over Na_2_SO_4_, and the solvent was evaporated. The residue was purified by column chromatography to afford ketone **S11**.

Br_2_ (1.1 equiv) was added dropwise to a stirred solution of **S11** (1.0 equiv) in 5.1 M HBr/HOAc (2.8 equiv) at room temperature. The reaction mixture was quenched with sat. Na_2_CO_3_ aq. and extracted three times with CH_2_Cl_2_. The combined organic layer was dried over Na_2_SO_4_, added TFA, filtrated, and concentrated *in vacuo* to give **S12** as the TFA salt.

To a solution of 2,4-dioxopiperidine-1-carboxylate (1.0 equiv) and **S12** (1.0 equiv) in EtOH (0.25 M) was added ammonium acetate (4.0 equiv) at room temperature. After stirring, the reaction mixture was quenched with sat. NaHCO_3_ aq., then the layers were separated. The aqueous layer was extracted with ethyl acetate and the combined organic layer was dried over Na_2_SO_4_ and concentrated *in vacuo*. The residue was purified by column chromatography to afford **S13**.

**S13** (10.0 mg, 1.0 equiv) in MeOH (0.3 mL) was added trimethylsilyl chloride (0.16 mL, 50 equiv) at room temperature. After stirring for 2 h, the reaction mixture was concentrated *in vacuo* to afford **AMI-138** and **AMI-215**.

**2-(2-Chloropyridin-4-yl)-1,5,6,7-tetrahydro-4*H*-pyrrolo[3,2-*c*]pyridin-4-one hydrochloride (AMI-138)**

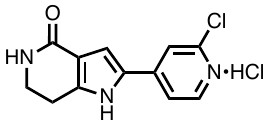

**AMI-138** was obtained as a yellow solid. ^1^H NMR (600 MHz, CD_3_OD) δ 8.27 (d, *J* = 5.4 Hz, 1H), 7.67 (s, 1H), 7.55 (dd, *J* = 5.4, 1.2 Hz, 1H), 7.13 (s, 1H), 3.61 (t, *J* = 7.2 Hz, 2H), 2.97 (t, *J* = 7.2 Hz, 2H); ^13^C NMR (150 MHz, CD_3_OD) δ 167.9, 148.5, 147.8, 146.6, 145.6, 129.2, 119.9, 118.5, 116.0, 112.9, 41.5, 22.7, one carbon signal was not detected even with prolonged scans; HRMS (ESI) *m/z* = 248.0585 calcd for C_12_H_11_ClN_3_O [M+H]^+^, found: 248.0582.

**2-(2,5-Dichloropyridin-4-yl)-1,5,6,7-tetrahydro-4H-pyrrolo[3,2-c]pyridin-4-one hydrochloride (AMI-215)**

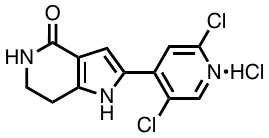

**AMI-215** was obtained as a yellow solid. ^1^H NMR (600 MHz, CD_3_OD) δ 8.38 (s, 1H), 7.66 (s, 1H), 7.40 (s, 1H), 3.58 (t, *J* = 7.2 Hz, 2H), 2.97 (t, *J* = 7.2 Hz, 2H); ^13^C NMR (150 MHz, CD_3_OD) δ 151.4, 151.0, 144.0, 141.0, 127.8, 122.6, 112.7, 41.7, 22.1, two carbon were overlapped with peaks, and amide carbon signal was not detected even with prolonged scans; HRMS (ESI) *m/z* = 282.0195 calcd for C_12_H_10_Cl_2_N_3_O [M+H]^+^, found: 282.0192.

**2.5 Synthesis of PHA hybrid–probe.**

**1-(11-Azidoundecyl)-3-bromo-2-(pyridin-4-yl)-1,5,6,7-tetrahydro-4*H*-pyrrolo[3,2-*c*]pyridin-4-one hydrochloride (AMI-327)**

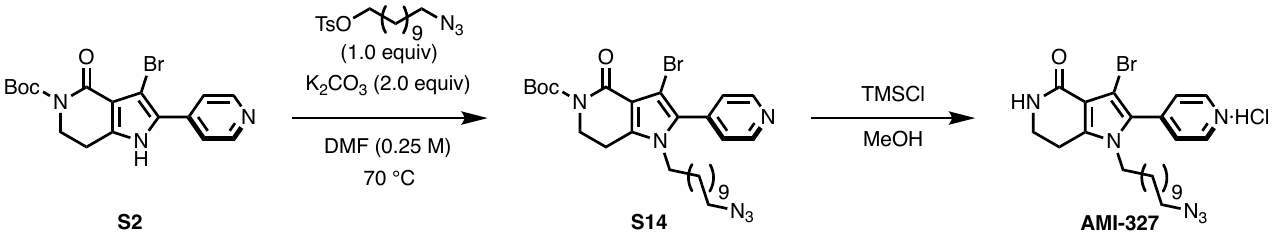

To a stirred solution of **S2** (695.0 mg, 1.77 mmol, 1.0 equiv) and K_2_CO_3_ (489.8 mg, 3.54 mmol, 2.0 equiv) in DMF (7 mL) was added 11-azidoundecyl 4-methylbenzenesulfonate (651.2 mg, 1.77 mmol, 1.0 equiv) at room temperature, and the mixture was stirred for an additional 23 h before being quenched by the addition of water. The reaction mixture was extracted with ethyl acetate, washed with brine, dried over Na_2_SO_4_, and concentrated *in vacuo*. The crude mixture was purified by flash column chromatography (CHCl_3_/MeOH = 100:0 to 19:1) to produce **S14** as a yellow oil.

**S14** (112.0 mg, 1.0 equiv) in MeOH (20 mL) was added trimethylsilyl chloride (1.2 mL, 50 equiv) at room temperature. After stirring for 2 h, the reaction mixture was concentrated *in vacuo* to afford **AMI-327** (39% yield). ^1^H NMR (600 MHz, CD_3_OD) δ 8.63 (d, *J* = 6.0 Hz, 2H), 7.47 (dd, *J* = 6.0, 1.2 Hz, 2H), 3.95 (t, *J* = 7.2 Hz, 2H), 3.52 (t, *J* = 7.2 Hz, 2H), 3.23 (t, *J* = 7.2 Hz, 2H), 2.93 (t, *J* = 7.2 Hz, 2H), 1.60–1.55 (m, 2H), 1.47–1.42 (m, 2H), 1.38–1.07 (m, 14H); ^13^C NMR (150 MHz, CD_3_OD) δ 167.0, 150.4, 141.2, 140.9, 130.9, 127.0, 112.7, 97.0, 52.4, 46.3, 41.1, 31.4, 30.5, 30.4, 30.3, 30.2, 29.9, 29.8, 27.8, 27.1, 22.8; HRMS (ESI) *m/z* = 509.1635 calcd for C_23_H_32_BrN_6_NaO [M+Na]^+^, found: 509.1638.

**1-(11-Azidoundecyl)-3-bromo-2-(pyridin-4-yl)-1,5,6,7-tetrahydro-4*H*-pyrrolo[3,2-*c*]pyridin-4-one (AMI-329)**

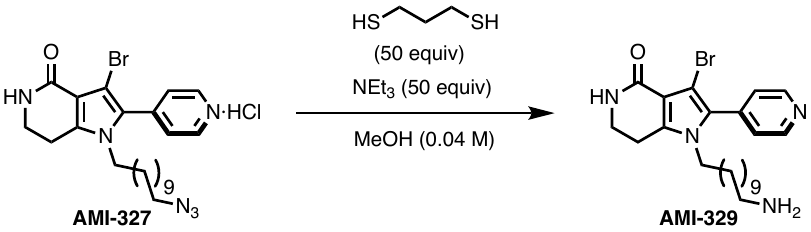

To a solution of **AMI-327** (64.0 mg, 0.13 mmol) and triethylamine (NEt_3_: 0.9 mL, 0.92 mmol, 50 equiv) in MeOH (20 mL) was added 1,3-propanedithiol (0.66 mL, 6.58 mmol, 50 equiv), and the mixture was stirred at room temperature for 39 h. The reaction mixture was concentrated*,* filtrated, and concentrated *in vacuo.* The residue was purified by PTLC (CHCl_3_/MeOH = 3:1, and 1% NEt_3_) to afford **AMI-329** as a yellow oil (75% yield). ^1^H NMR (600 MHz, CD_3_OD) δ 8.64 (dd, *J* = 6.0, 1.2 Hz, 2H), 7.50 (dd, *J* = 6.0, 1.2 Hz, 2H), 3.99 (t, *J* = 7.2 Hz, 2H), 3.56 (t, *J* = 7.2 Hz, 2H), 2.96 (t, *J* = 7.2 Hz, 2H), 2.76 (t, *J* = 7.2 Hz, 2H), 1.57–1.52 (m, 2H), 1.46–1.40 (m, 2H), 1.36–1.03 (m, 14H); ^13^C NMR (150 MHz, CD_3_OD) δ 167.0, 150.5, 141.1, 140.9, 130.9, 127.0, 112.7, 96.9, 49.9, 46.3, 41.7, 41.1, 31.3, 30.5, 30.4, 30.4, 30.3, 29.7, 27.7, 27.1, 22.8; HRMS (ESI) *m/z* = 461.1911 calcd for C_23_H_34_BrN_4_O [M+H]^+^, found: 461.1911.
